# A Statistical Method to Estimate the Population-Level Frequencies of *Plasmodium falciparum* Haplotypes with *Pfhrp*2/3 Deletions in the Presence of Mixed-Clone Infections

**DOI:** 10.64898/2026.04.01.715806

**Authors:** Loyce Kayanula, Kanika Verma, Praveen Kumar Bharti, Kristan Alexander Schneider

## Abstract

**Background:** The World Health Organization (WHO) has raised concerns over increasing *Pfhrp*2/3 deletions, undermining the sensitivity of *Pfhrp*2-based rapid diagnostic tests (RDTs). Close monitoring of the population and a change in diagnostic methods are recommended if the prevalence of parasites with *Pfhrp*2/3 deletions exceeds 5%. In high transmission settings, accurate estimates are hampered by the frequent occurrence of mixed-clone infections (multiplicity of infection; MOI).

**Objective and Methods:** If parasites with and without deletions are present in an infection, standard molecular assays cannot detect the presence of the former. To accurately estimate frequencies of haplotypes with *Pfhrp*2/3 deletions in the presence of mixed infections, a novel statistical model that combines genetic/molecular information from *Pfhrp*2/3 with that from neutral markers is introduced. Maximum-likelihood estimates (MLEs) are obtained for haplotype frequencies characterized by markers at *Phrp*2/3 loci and loci for neutral markers. The expectation-maximization algorithm is used to derive the MLEs. The adequacy of the method (precision and accuracy) is assessed by numerical simulations.

**Results:** The method was applied to an active surveillance study conducted in a tribal community in Jagdalpur, India, which enrolled febrile community members (*n* = 432) between October and November 2021. Four markers each at *Pfhrp*2 and *Pfhrp*3 are combined with one marker each at *Pfmsp*1 (which encodes *P. falciparum* merozoite surface protein 1) and *Pfmsp*2. Data from a total of 117 patients who had both *P. falciparum* infections and genetic information for the molecular markers underwent further analysis with the novel statistical method.

**Conclusion:** Results indicate that this novel method has promising statistical properties (asymptotic and in finite samples) and can be readily applied to real-world situations. A stable implementation of the method in R is provided. This novel approach enables accurate estimation of *Pfhrp*2/3 deletion frequencies in complex *P. falciparum* infections, addressing a key limitation of current molecular surveillance methods.

**Author summary:** *Plasmodium falciparum* (*Pf*) causes the most severe form of human malaria, accounting for over 90% of cases. Rapid diagnostic tests (RDTs) have become a cornerstone of malaria control. These RDTs detect *Pf*-specific antigens in a blood drop. HRP2/3 emerged as the best antigen for such tests because it is *Pf*-specific and expressed in abundance. However, some parasites lack the genes that code for HRP2/3 proteins. If parasites in an infection have such gene deletions, RDT results can be false negative. The WHO considers the containment of such deletions a public health priority and recommends monitoring their prevalence. The detection of HRP deletions is challenging if parasites with and without deletions co-occur in infections because standard molecular assays cannot detect deletions in this situation. To overcome this challenge, we introduce a novel statistical method to estimate the frequency distribution of parasite variants with deletions. The method combines information from neutral molecular markers and from HRP-related markers to correct for unobservable information. Here we provide a derivation of the statistical model, a stable implementation, and test its statistical properties with synthetic and real data, thereby showing that our method is well-suited for the underlying problem.

## Introduction

Despite a reduction in estimated malaria cases per population at risk over the past 25 years, the disease still caused 282 million estimated cases and 0.6 million deaths in 2025 [1]. The majority of infections are caused by *Plasmodium falciparum* (*P. falciparum*), the clinically most relevant malaria species. After years of decline, the number of cases and deaths per population at risk has been increasing again since 2018. The number of confirmed malaria infections has been steadily increasing over the past quarter century, mainly attributed to better diagnostics.

Historically, malaria diagnosis relied on unspecific clinical symptoms, such as fever and headache, which overlap with many other common febrile illnesses [2, 3]. This led to widespread misuse of antimalarial drugs and facilitated the emergence of drug resistance [4, 5]. With increased awareness of drug resistance, the global shift toward artemisinin-based combination therapies (ACTs) to treat *P. falciparum* malaria was accompanied by the introduction of disease-specific diagnostics. Although light microscopy remains the “gold standard” for parasite detection and species identification, it is often unavailable in resource-limited settings due to requirements for skilled personnel, equipment, and time [6, 7]. The roll-out of rapid diagnostic tests (RDTs) since the early 2000s mitigated this problem. These antigen-based lateral-flow chromatography tests have the same sensitivity as light microscopy and have improved malaria diagnosis by allowing infections to be confirmed quickly without the need for a laboratory infrastructure.

The most common RDTs target either the *P. falciparum* specific histidine-rich protein 2 (HRP2) or the pan-species parasite lactate dehydrogenase (pLDH). An HRP2-based test confirms the presence of *P. falciparum*, the species for which ACTs are first line therapy, whereas a pLDH-based test indicates a malaria infection but cannot by itself specify the species [8, 9]. Consequently, HRP2-based RDTs, which also detect the structurally similar HRP3, are recommended by the WHO before treatment (especially in *P. falciparum* endemic regions). Notably, some RDTs target HRP2 and HRP3. RDTs also offer a step of verification for infection with low density parasitemia, which were negative by light microscopy, especially at field sites [7, 9]. Moreover, they are particularly valuable in specific patient groups, such as those with sickle cell disease (SCD), where low parasite densities can be easily missed by microscopy [10].

The performance of HRP2-based RDTs depends on the expression of their target antigens, HRP2 and HRP3, which are encoded by the *P. falciparum* histidine-rich protein 2 and 3 (*Pfhrp*2/3) genes [11–14]. However, molecular surveillance studies have revealed the emergence and spread of parasite haplotypes with deletions in these genes. These deletions, which can be complete (removing the entire gene) or partial, were first documented in parasite isolates from the Peruvian Amazon [15] and have since been reported in multiple malaria endemic regions worldwide [12, 16, 17]. By eliminating the target antigens, these deletions compromise the accuracy of HRP2-based RDTs, leading to false negative results and subsequent treatment delays or inappropriate medication [15–17]. The spread of *Pfhrp*2/3 deletions directly threatens the strategy pursued by global health authorities to confirm suspected malaria cases by RDT or microscopy before treatment (“test-and-treat” strategy) [18, 19]. Furthermore, evidence suggests that parasites carrying *Pfhrp*2/3 deletions may be more likely to cause asymptomatic infections [2, 20], which naturally evade diagnosis and treatment.

Due to their importance, the WHO recommends monitoring *Pfhrp*2/3 deletions and proposes a critical threshold of 5% prevalence of *Pfhrp*2/3 deletions among symptomatic patients as an indicator for switching to non-HRP2-based RDTs [17, 21].

In high transmission settings, individuals are often super-infected with multiple genetically distinct parasite clones during a single disease episode, a phenomenon known as multiplicity of infection (MOI) [22–24]. If wild-type haplotypes and those having deletions co-occur in mixed infections, the presence of functional alleles from the wild types mask the co-existence of deletions in molecular assays (see last row “Observations” fourth column in Fig 2) [21]. This ‘masking effect’ likely leads to a substantial underestimation of the true prevalence of *Pfhrp*2/3 deletions in parasite populations in high-transmission settings.

Current statistical methods for estimating haplotype frequencies and MOI are not designed to estimate frequencies of *Pfhrp*2/3 deletion haplotypes. Existing approaches typically treat absent allelic information at a marker as missing data rather than a biological indicator of gene deletion [25]. Although methods exist to explicitly include missing information, e.g., [26], these frameworks regard missing data as a consequence of laboratory failure rather than systematic allele absence caused by gene deletions. In the context of *Pfhrp*2/3 deletions, their presence in mixed infections, which render them unobservable, is more important. Current methods cannot accurately estimate *Pfhrp*2/3 deletion haplotype frequencies or correct for biases because they can be unobservable in polyclonal infections.

To address these limitations, a novel statistical framework that simultaneously estimates frequencies of haplotypes with *Pfhrp*2/3 deletions and MOI is presented. This method combines markers at which deletions can occur with neutral genetic markers. The latter are included to create enough resolution to detect polyclonal infections and overcome the intrinsic problem of deletions being unobservable if co-occurring with variants that do not have deletions.

Here, a formal statistical model to estimate the frequency distribution of haplotypes having *Pfhrp*2/3 deletions is introduced. The likelihood function is derived and the Expectation-Maximization (EM) algorithm [27] is used to numerically obtain the maximum-likelihood estimates. Within this statistical framework, formulae to calculate the prevalence of *Pfhrp*2/3 deletions (i.e., the probability that these are present in an infection) are derived. The asymptotic variance of the estimator in terms of the inverse Fisher information is derived and it is shown that the variance of the estimator is in good agreement with the Cramér-Rao lower bound to establish the theoretical efficiency of the estimators [28]. Through simulations, it is demonstrated that the method presented provides accurate and precise estimates for haplotype frequencies and MOI across various a range of parameters and sample sizes. Finally, the method is applied to empirical data from a study conducted in a tribal community in Jagdalpur, India. The method is implemented as an R script available as supplementary material and at GitHub https://github.com/Maths-against-Malaria/Estimating-HRP-deletions.

This approach provides the necessary tools to accurately monitor *Pfhrp*2/3 deletions in accordance with WHO recommendations in areas of high transmission, in which polyclonal infections are common and lead to underestimates of the frequency of such deletions.

## Results

This section presents the results based on the statistical model derived in Methods. First, the algorithm for obtaining the maximum likelihood estimates (MLEs) of the *Pfhrp*2/3 deletion haplotype frequencies and the MOI parameter under the proposed model is described. Second, *Pfhrp*2/3 deletion haplotype prevalence estimates are derived using the MLEs as plug-in estimates. Third, the asymptotic variance of the estimator is derived via the Fisher information, yielding the corresponding Cramér–Rao lower bound. Finally, the finite sample performance of the estimator is investigated through numerical simulations.

The results are presented using specific notations intended to improve clarity and to support the derivations described in Methods. Table 1 summarizes the critical notations.

**Table 1.**
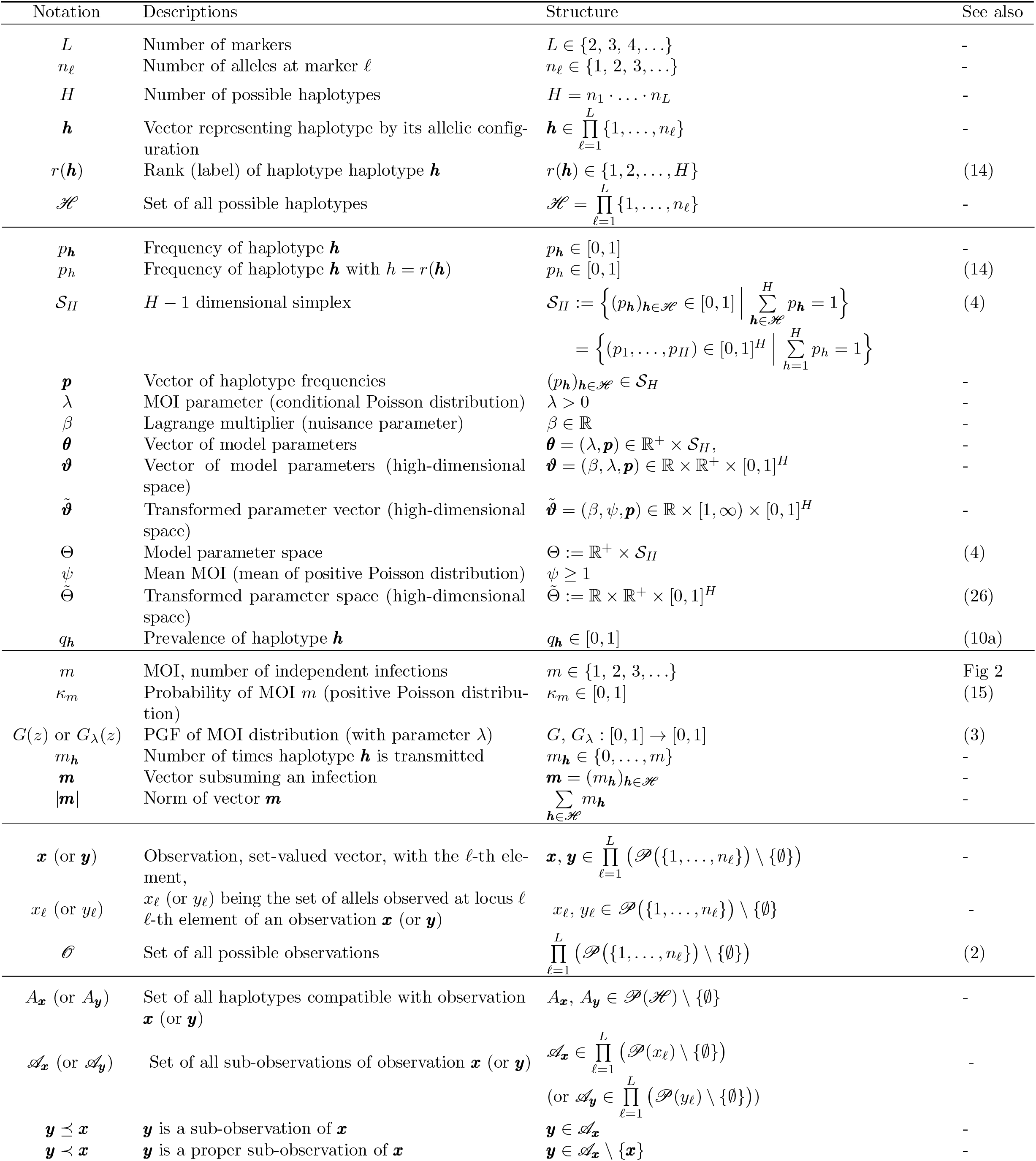
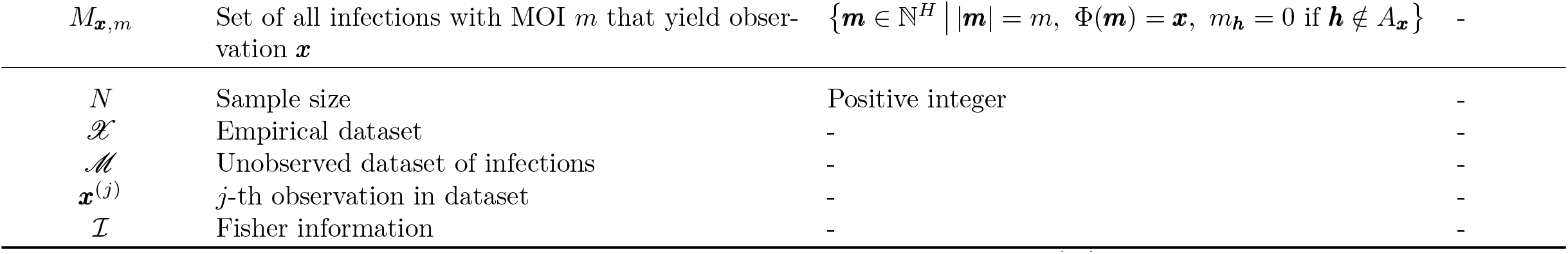
Summary of key notation: *𝒮*_*H*_ denotes the (*H* − 1)-dimensional simplex, 𝒫 (*X*) is the powerset of *X*, and PGF stands for probability generating function.

### Model overview

A concise derivation of the statistical model is given in Methods. Here, the model is briefly summarized to facilitate readability of the results. The key notation is summarized in Table 1.

The probability distribution of observations that are obtained from molecular assays concerning *Pfhrp*2/3 deletions is modeled. For this purpose a genetic architecture of *L* multi-alellic markers (loci) is considered. The *n*_*ℓ*_ alleles segregating at marker *ℓ* are denoted by 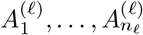. The first *D* markers correspond to regions in *Pfhrp*2/3. The convention is used that the first allele 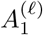 at loci 1, …, *D* corresponds to a deletion, while the remaining alleles are distinct variants.

With this genetic architecture in mind, a parasite haplotype ***h*** is characterized by specific allelic variants at the *L* markers. Formally, a haplotype is denoted as a vector ***h*** = (*h*_1_, …, *h*_*L*_), where *h*_*ℓ*_ ∈ {1, …, *n*_*ℓ*_}. The set of all 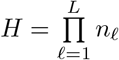 possible haplotypes is denoted by 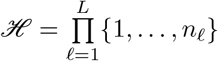.

A haplotype ***h*** with deletions at all of the first *D* markers is referred to as “complete deletion”. A haplotype with deletions at some but not all of the first *D* markers is referred to as “partial deletion”. The advantage of partitioning the genetic architecture at *Pfhrp*2/3 into *D* markers is that partial deletions can be easily modeled (cf. Fig 1).

**Fig 1.**
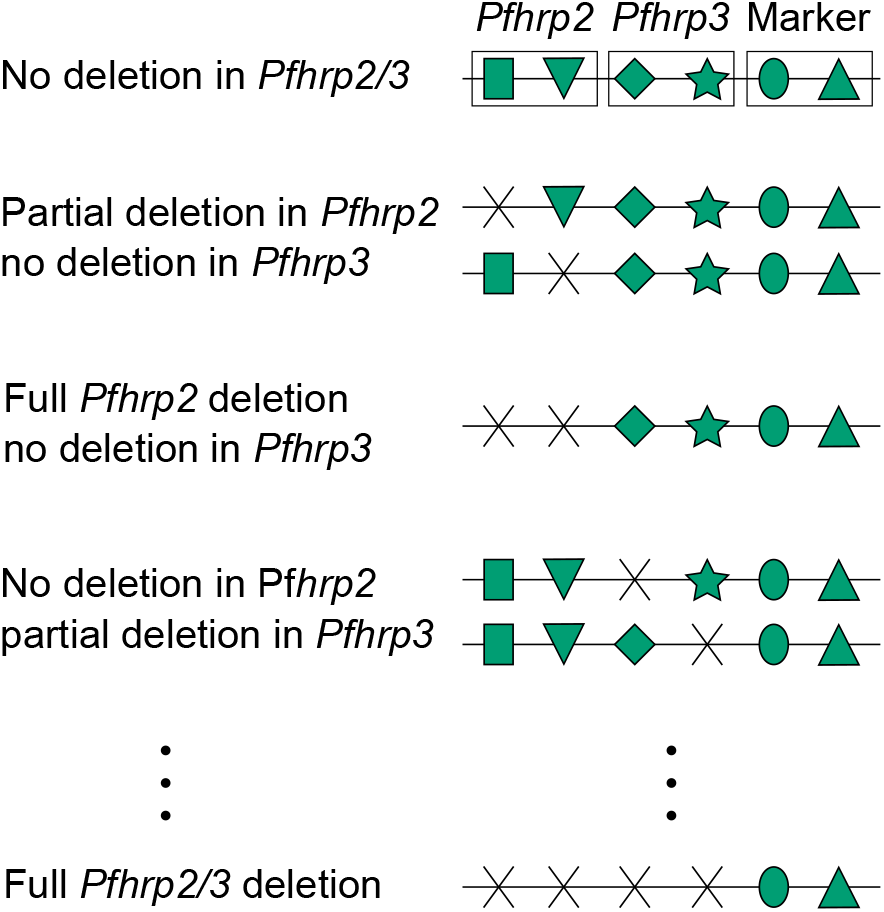
Illustration of genetic architecture: Illustrated is the genetic architecture assuming two markers in *Pfhrp*2, two markers in *Pfhrp*3 and two additional markers. Each horizontal line illustrates the genome of a parasite haplotype. The symbols illustrate different alleles. Here, deletions are illustrated by an ‘X’. Illustrated are seven haplotypes, with different *Pfhrp*2/3 deletions.

The sporozoite stage in the mosquito salivary glands serves as the census point for the haplotype frequency distribution [22], which are denoted by the vector ***p*** := (*p*_***h***_)_***h***∈ℋ_.

Parasites are observed in human hosts during the erythrocytic cycle. A confounding factor is that hosts can be super-infected due to multiple disease exposures. Here, super-infections refer to independent infective events during one disease episode. It is further assumed that at each infecting event, exactly one parasite haplotype is transmitted to the host. This ignores the possibility of co-infections, i.e., the transmission of multiple parasite haplotypes during one infective event (for more discussion on this assumption, see e.g. [29]). Following [22, 24, 30] the number of super-infections is referred to as multiplicity of infection (MOI).

Assuming infections are rare and independent, the number of infections per individual, denoted by *m*, is a realization of a random variable following a Poisson distribution. Considering only disease-positive hosts, MOI follows a conditional or positive Poisson distribution (*m* ~ CPoiss(*λ*)). The mean MOI, denoted by *ψ*, is calculated to be

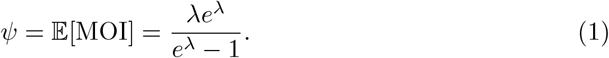

Notably, the derivation of the statistical model and the likelihood function in the following are independent of the assumption of the Poisson distribution and can be changed by any other distribution of the positive integers. However, derivation of the EM algorithm, is based on the Poisson assumption. In the following, the probability generating function of the MOI distribution is denoted as *G*, or *G*_*λ*_ in case it is necessary to indicate that the positive Poisson distribution is assumed (see Methods for details).

The assumption of exactly one parasite haplotype being transmitted at each infectious event, implies that, given MOI *m*, the configuration of haplotypes infecting a host follows a multinomial distribution with parameters *m* and ***p***. Given MOI = *m*, let *m*_***h***_ denote the number of times the host was infected with haplotype ***h*** (subject to the constraint 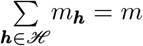).

In practice, MOI is unobservable and haplotype information is typically unavailable because molecular assays yield unphased information (cf. Fig 2). More precisely, at each marker *ℓ* the set of alleles *x*_*ℓ*_ ⊆ {1, …, *n*_*ℓ*_} present in the infection is observable, it is in general unknown which haplotypes are present in an infection (i.e., for which haplotypes ***h*** the number *m*_***h***_ *>* 0). As the first allele 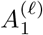 corresponds to a deletion at markers *ℓ* = 1, …, *D*, which is by nature “unobservable”, the convention 1 ∈ *x*_*ℓ*_ for *ℓ* = 1, …, *D* is made here. Formally, in a clinical sample at marker *ℓ* the observable information is

**Fig 2.**
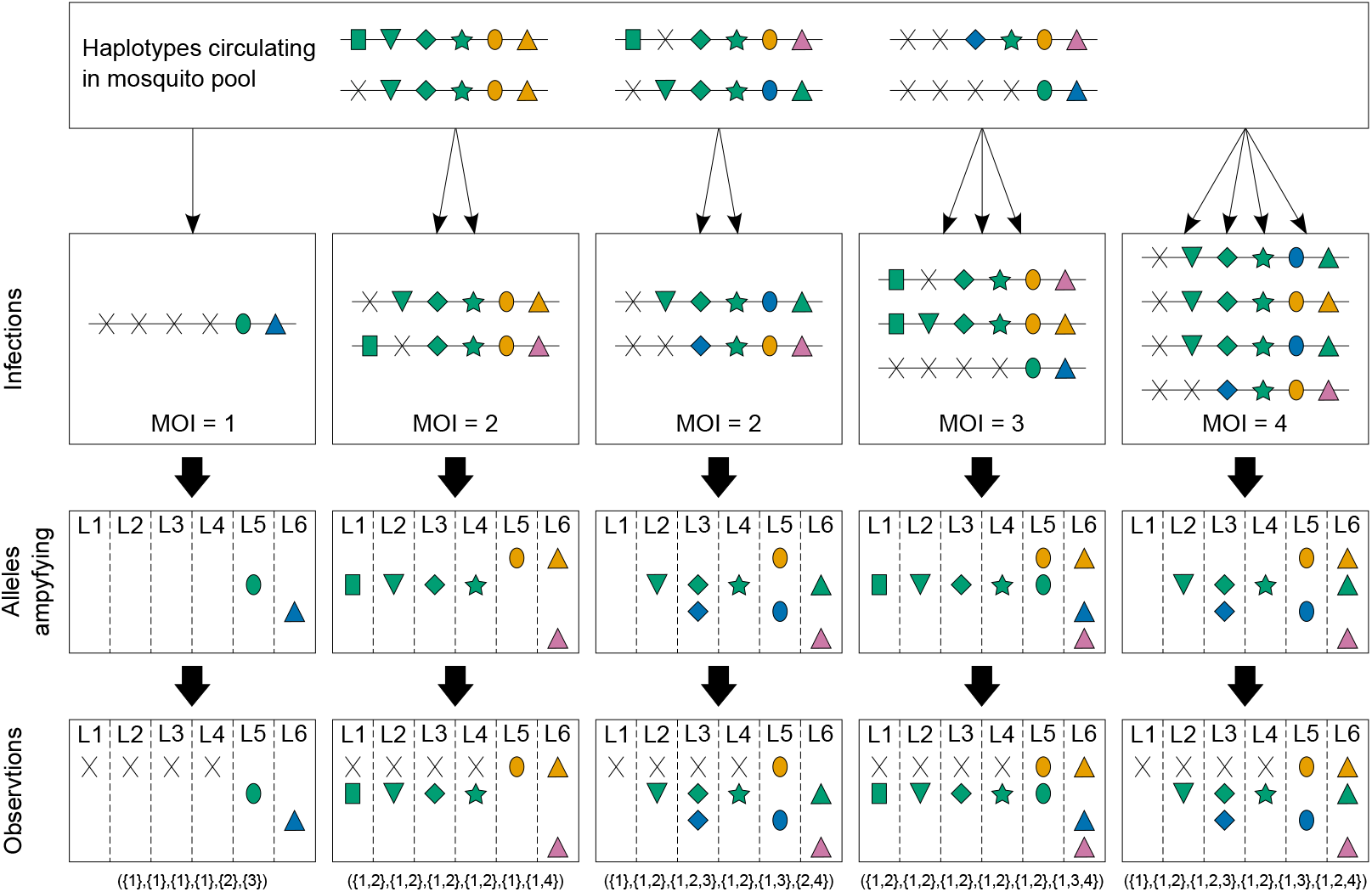
Illustration of infections and observations: Illustrated is a parasite population with six circulating haplotypes (top row) and five possible infections (second row). The haplotypes are illustrated as in 1, but here, different colors indicate different alleles. The five infections have varying MOI. Note that the last infection has MOI = 4, but only three haplotypes are transmitted since one haplotype is infecting twice. In all infections, at least one parasite haplotype is present with a deletion. The third row shows the resulting alleles which are detected by a molecular/genetic assay. Note, if all haplotypes in an infection have a deletion at the same position, no allele information is obtained (see loci L1-4 in the first infection, and marker L1 in the third and fifth infection). In the second and fourth infection, allelic information is obtained for all markers. Although parasites with deletions are present in all five infections, due to multiple infections, this cannot be seen from the allelic information of the second and fourth infection. Moreover, from the allelic information in the third and fifth infections, it is only apparent that a deletion occurs at the first position in *Pfhrp*2. Since deletion will result in null-information in molecular assays, the resulting ambiguity needs to be considered in the observations (bottom row). At the bottom of each panel, the representation of the samples as elements in 𝒪 is shown.

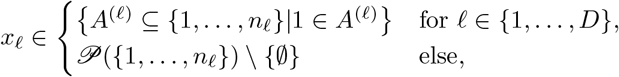

with 𝒫 denoting the power set. (Note, the empty set is excluded because only disease positive samples are considered.) The set of all possible observations is

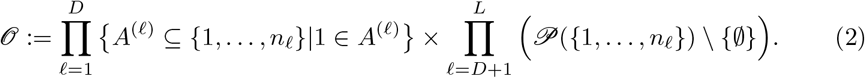

An observation ***y*** a called a sub-observation of ***x***, denoted ***y*** ⪯ ***x***, if all alleles observed in ***y*** are also observed in ***x*** (see Methods for a formal description). The set of all sub-observations of ***x*** is defined by

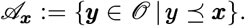

Because, in general, it is unknown which haplotypes are present in an infection. However, from an observation ***x*** the presence of certain haplotypes in the infection can be ruled out, as these are incompatible with the observation ***x***. The set of all haplotypes, “compatible” with observation ***x***, is denoted by

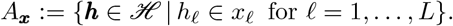

Note, if a haplotype ***h*** is an element of *A*_***x***_ it does not imply that haplotype ***h*** was present in the infection leading to observation ***x***, rather its presence cannot be ruled out.

The probability observing ***x*** is derived in detail in Methods and given by

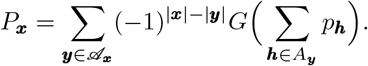

When assuming that MOI follows a conditional Poisson distribution, then the PGF is given by

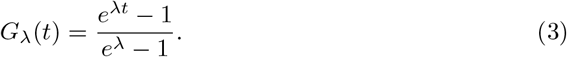

In this case, the parameter space of the model is a *H*-dimensional space Θ defined as

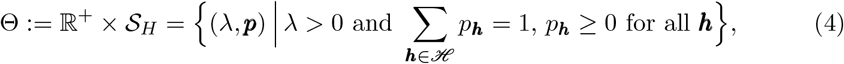

where 𝒮_*H*_ denotes the *H* − 1-dimensional simplex.

### Maximum likelihood estimation

Assume a dataset 𝒳 consisting of *N* observations. The *j*-th observation is denoted by 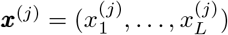, where 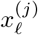 is the allelic information at marker *ℓ* in sample *j*. The number of times observation ***x*** occurs in the dataset 𝒳 is denoted by *n*_***x***_, such that 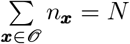.

Given 𝒳, the model parameters ***θ*** := (*λ*, ***p***) ∈ Θ can be collectively estimated by the maximum-likelihood method. The log-likelihood function (see Methods) is

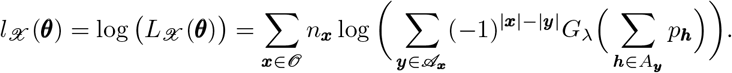

This log-likelihood function does not permit a closed-form solution for the maximum-likelihood estimate (MLE). The estimates of the *Pfhrp*2/3 haplotype frequencies ***p*** and the MOI parameter *λ* are hence obtained by numerical maximization using the expectation-maximization (EM) algorithm.

The EM algorithm is an iterative method composed of two alternating steps. The expectation (E)-step computes the conditional expectation of the log-likelihood, given the observed data and current parameter estimates, with respect to the unobserved variables. The maximization (M)-step updates the parameter estimates by maximizing this expected log-likelihood. These steps are repeated until a predefined convergence criterion is met, yielding the maximum likelihood estimates (MLEs). A detailed derivation of the algorithm is provided in section Maximizing the likelihood function with the Expectation-Maximization (EM) algorithm in Methods.

Given the number of times an observation ***x*** is observed in a dataset denoted as *n*_***x***_, the algorithm begins with initial values ***p***^(0)^ and *λ*^(0)^. At iteration *t*, given current estimates ***p***^(*t*)^ and *λ*^(*t*)^, the updates for step *t* + 1 are computed as follows. The *Pfhrp*2/3 haplotype frequencies are updated according to

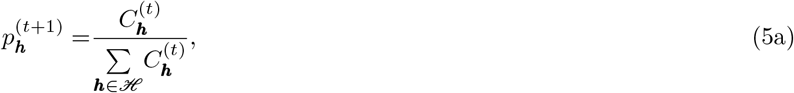

where

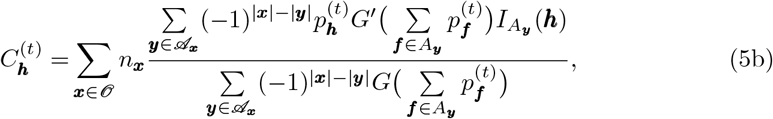

with

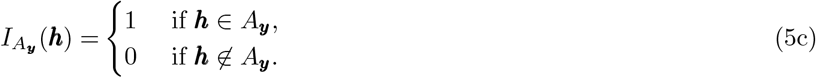

After updating the haplotype frequencies, the MOI parameter is updated by solving the following fixed-point iteration

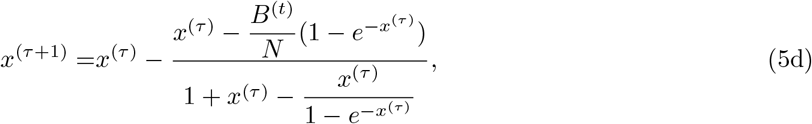

with

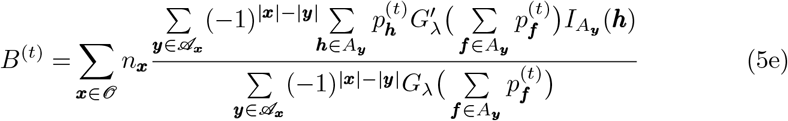

Starting from *x*^(0)^ = *λ*^(*t*)^, (5d) is iterated until convergence. The resulting converged value *x*^(*τ*+1)^ is then taken as the updated MOI estimate, i.e., *λ*^(*t*+1)^. The two-step iteration above is repeated until convergence, i.e., until 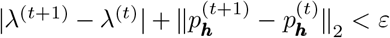, for a given threashold *ε*. From 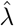, the mean MOI *ψ* is estimated as

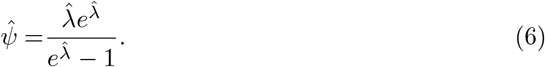

The EM algorithm is guaranteed to converge to a local maximum of the likelihood. In practice, convergence is typically achieved within a modest number of iterations. See Table 1 for all the symbols and notations used. An implementation of the algorithm is available via an R script that can be found at GitHub https://github.com/Maths-against-Malaria/Estimating-HRP-deletions.

### Fisher Information and the Cramér-Rao Lower Bound

The precision of parameter estimators can be quantified using the Cramér-Rao lower bound (CRLB), which is the inverse Fisher information and provides a theoretical minimum variance achievable by an unbiased estimator [31, 32]. Maximum-likelihood estimators are often only asymptotically unbiased, so this lower bound does not apply. However, in practice, if an asymptotically unbiased estimator has a variance close to the CRLB there is little hope for improvement.

For any unbiased estimator Φ for the true parameter vector ***θ***_0_ ∈ Θ

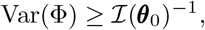

where ℐ (***θ***_0_) is the Fisher information matrix.

In the present case, the parameter space Θ is constrained, by the haplotype frequencies ***p*** satisfying 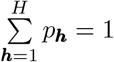. The method of Lagrange multipliers can again be used to to derive the inverse Fisher information following the arguments in [25], which uses a higher-dimensional embedding with a Lagrange multiplier *β* as nuisance parameter, i.e., the higher-dimensional space defined in 26 is used.

Using the extended parameter vector 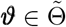 defined in (27), we have the following Lagrangian function

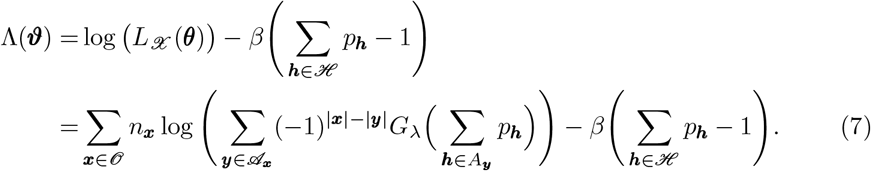

The Fisher Information matrix for ***ϑ*** is then defined as

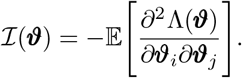

After deriving the necessary first and second order derivatives (see Fisher Information in S1 Appendix), the components of ℐ (***ϑ***) are

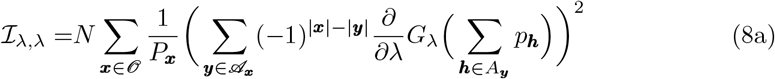

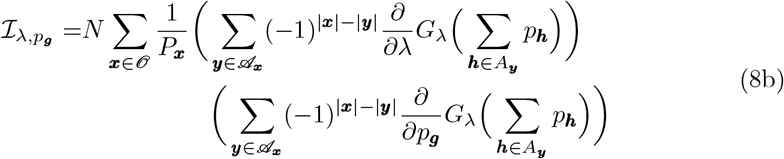

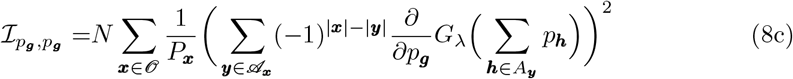

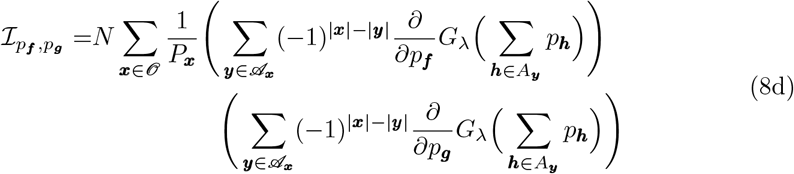

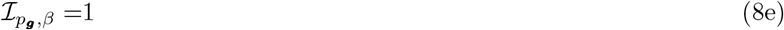

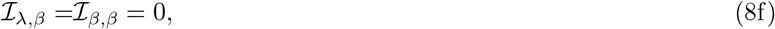

where

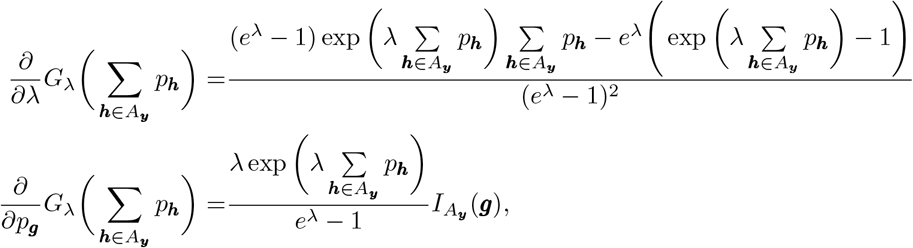

and 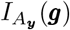 is as defined in (24). The resulting Fisher Information matrix has the block structure

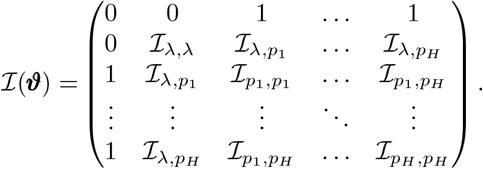

The inverse matrix ℐ (***ϑ***)^−1^, disregarding entries corresponding to the nuisance parameter *β* (i.e., first row and column), is the CRLB for the parameters of interest, ***θ***, with the diagonal entries representing minimum achievable variances and off-diagonal entries representing covariances between estimates.

Recall, under the Conditional Poisson distribution, the mean MOI *ψ* is related to *λ* via

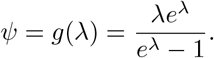

The Lagrangian function can be expressed in terms of *ψ* as

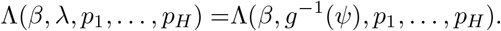

This formulation allows the Fisher information to be computed directly for 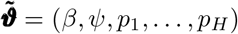. Using the chain rule, the first and second order derivatives for this Lagrangian function are obtained as shown in section Fisher Information in S1 Appendix). The corresponding entries for the Fisher information are as follows

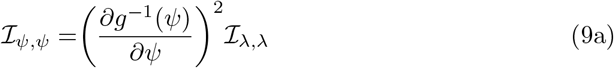

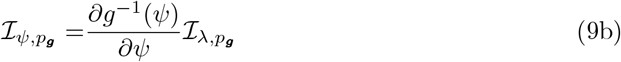

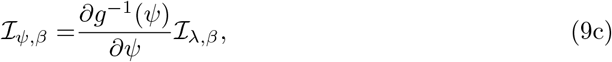

with 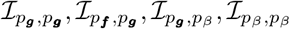 as obtained in (8) above. Therefore, using (8), and (9), the Fisher Information matrix for the parameters 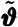 can be written as

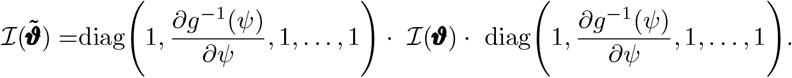

Hence,

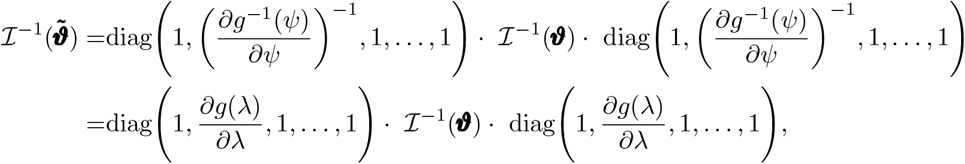

where

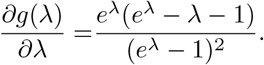

The Fisher information and the associated Cramér–Rao lower bounds are thus obtained for the reparameterized model.

### Observed and unobserved deletions within infections

The frequency of haplotype ***h*** is its relative abundance in the parasite population; its prevalence is the probability that haplotype ***h*** is present in an infection. If exactly one haplotype was infecting, prevalence and frequency would coincide. However, due to multiplicity of infection, haplotype prevalence exceeds frequency [33].

In the context of HRP deletions, it is important to ascertain how many infections will yield a false-negative RDT result due to lack of HRP antigens. Hence, the probability of infections in which all parasites have deletions at one or several positions in the *Pfhrp*2/3 are of interest.

#### Prevalence

The prevalence of haplotype ***h*** is denoted by *q*_***h***_. The expression follows readily from the counter probability of haplotype ***h*** not being present in an infection. Namely,

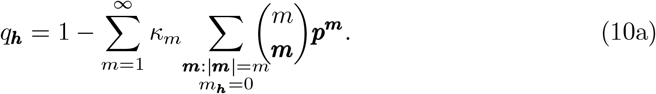

The multinomial theorem gives

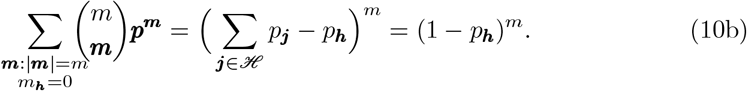

Therefore,

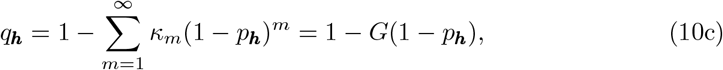

where again *G* is the probability generating function of the MOI distribution. (Note, this argument is general, and does not require that MOI is conditionally Poisson distributed.)

#### Infections without fully functional HRP

If some haplotypes in an infection have a deletion at a given position, while some others do not, the deletion is not directly observable. Potentially, the sensitivity of RDTs is reduced even in infections without directly observable deletions. Namely, assume two haplotypes are super-infecting, the first having a deletion at position 1, the second at position 2. If the ability to produce functional HRP is restricted in both haplotypes, no fully functional HRP will be circulating in the infection; however, the deletions are not observable. It is important to determine the proportion of infections for which the sensitivity of RDTs is reduced. There are several cases of interest that fall into two situations. First, the cases in which deletions are directly observable in infections, and second cases in which deletions are not observable, yet no fully functional HRP is circulating in the infection.

Recall that the first *D* of the *L* considered markers correspond to positions in *Pfhrp*2/3. Let *A* ⊆ {1, …, *D*} =: 𝒟 be a set of positions in the *Pfhrp* genes, and define the set of haplotypes that have deletions at all positions in *A* as

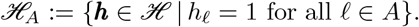

A set of deletions *A* ⊆ 𝒟 is directly observable in an infection if all parasites in that infection have haplotypes in ℋ_*A*_, i.e., all infecting haplotypes carry deletions at all positions in *A*. The prevalence of such infections (i.e., the probability that an infection consists entirely of haplotypes from ℋ_*A*_), is denoted as *q*_*A*_. Then

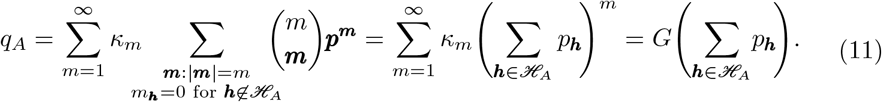

If *A* = {*ℓ*} for some *ℓ* ∈ {1, …, *D}*, this gives the prevalence of infections in which all parasites have a deletion at position *ℓ*.

Now consider infections in which no fully functional HRP is produced, yet the deletions causing the loss of function are only partially observable, or not observable at all, due to their distribution across different haplotypes. Define the set of haplotypes that have no deletion at any position not in *B* as

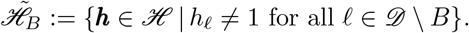

A set of deletions *A* ⊆ *B* is directly observable in an infection if all infecting haplotypes belong to ℋ_*A*_. Next, define

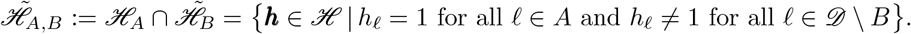

If only haplotypes in 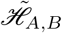 are present in an infection, deletions at positions in *A* are directly observable while no deletion outside *B* is present. The probability of infections containing only haplotypes in 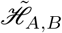 is

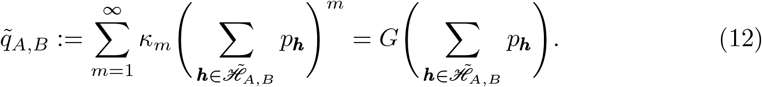

The deletions at positions in *A* are directly observable, while it is unclear whether some are present at positions in *B \A*. Therefore, the probability that the set of haplotypes present in an infection satisfies that no haplotype has a deletion at a position outside *B*, all have deletions at all positions in *A*, and for every position in *B \A* at least one haplotype has a deletion is – by using an inclusion-exclusion agrument – given by

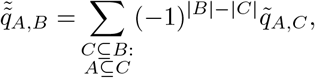

where 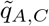 is given by (12). Using another inclusion-exclusion argument yields the probability of observations in which deletions are present at exactly the positions in *B*, and are directly observable at exactly the positions in *A*. Namely,

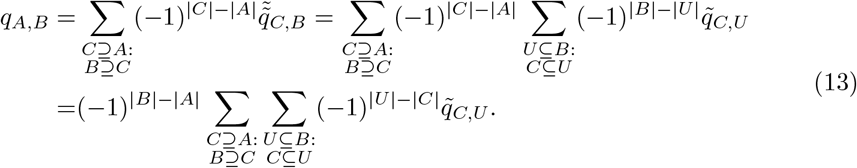

If *A* = *B*, then *B \A* = ∅ and (13) reduces to *q*_*B*_, the prevalence of directly observable deletion patterns derived in (11). Of particular interest is the case *A* = ∅, i.e., the probability of infections in which all haplotypes have deletions at one or more positions in *B*, but no deletion is directly observable. In such infections, no functional HRP is produced, affecting the sensitivity of RDTs, but deletions are unobservable.

### Finite sample properties

The performance of the estimator was assessed under varying conditions, including different sample sizes, initial haplotype frequencies, and transmission levels (see Figs 3,4, 5). The theoretical efficiency of the estimator was validated by comparing empirical precision with the Cramér-Rao lower bound derived from the Fisher Information. The objective of the estimator is to correctly estimate the frequencies of *Pfhrp*2/3 deletions. Hence, the first *D* markers are of primary interest. Markers *D* + 1, …, *L* have to be included in the method, but haplotype information involving these markers is not of interest. In other words, of primary interest are the frequency estimates marginalized to the first *D* markers. Hence, all results on frequencies presented here are marginalized.

**Fig 3.**
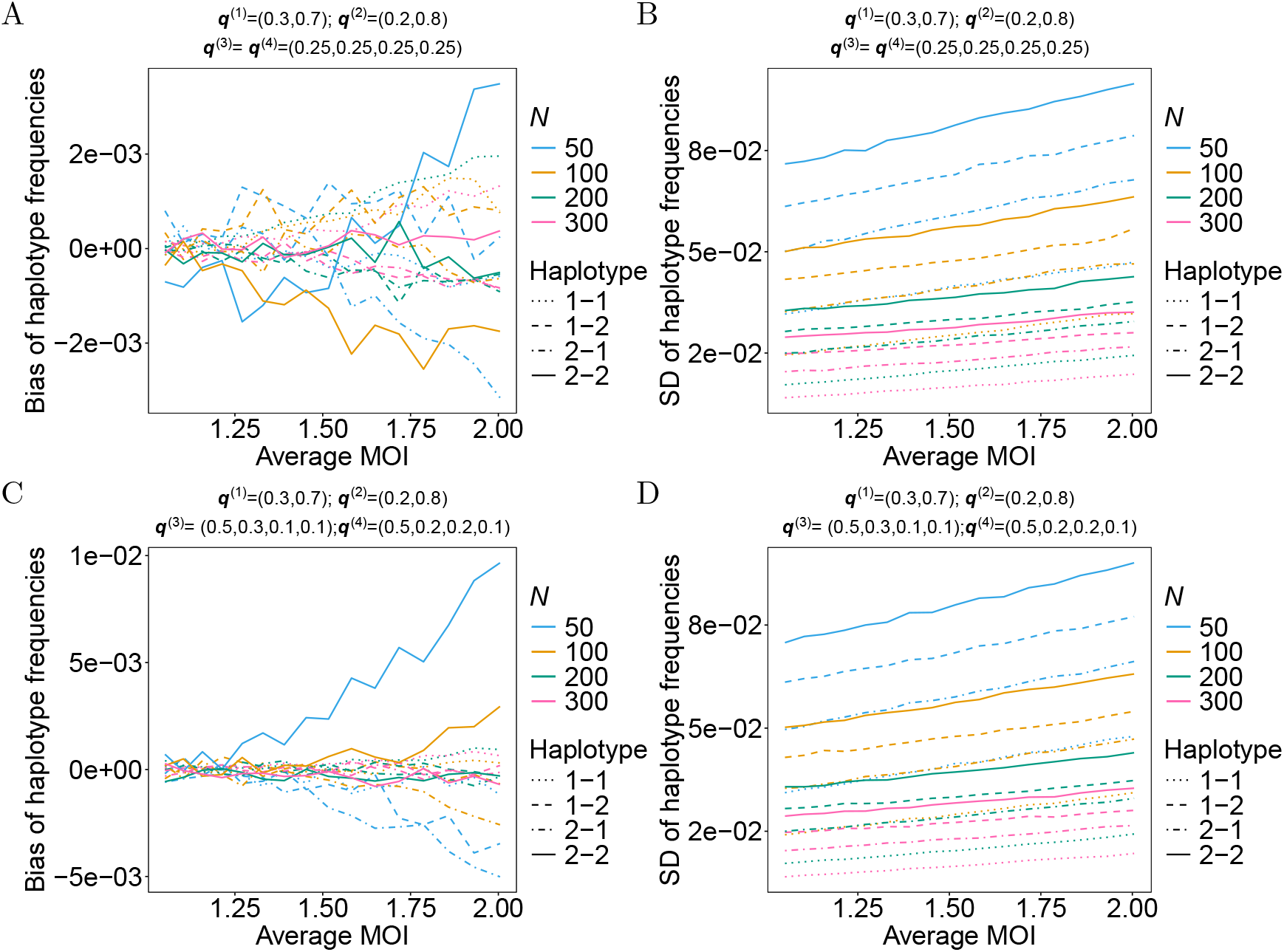
Bias and standard deviation of haplotype frequency estimates as functions of verage MOI. Panels A and C show the bias of haplotype frequency estimates as a function of average MOI for different sample sizes (*N*; colors), while panels B and D show the standard deviation (SD). True underlying haplotype frequencies were derived from fixed allele frequencies (shown at the top of each panel) under the assumption of linkage equilibrium. Results for the four resulting marginalized haplotypes are shown by different line types. Line types represent the four different haplotype deletion types. In panels A and B true allele frequencies at neutral loci (*ℓ* = 3, 4) are uniform. In panels C and D true allele frequencies are unbalanced.

**Fig 4.**
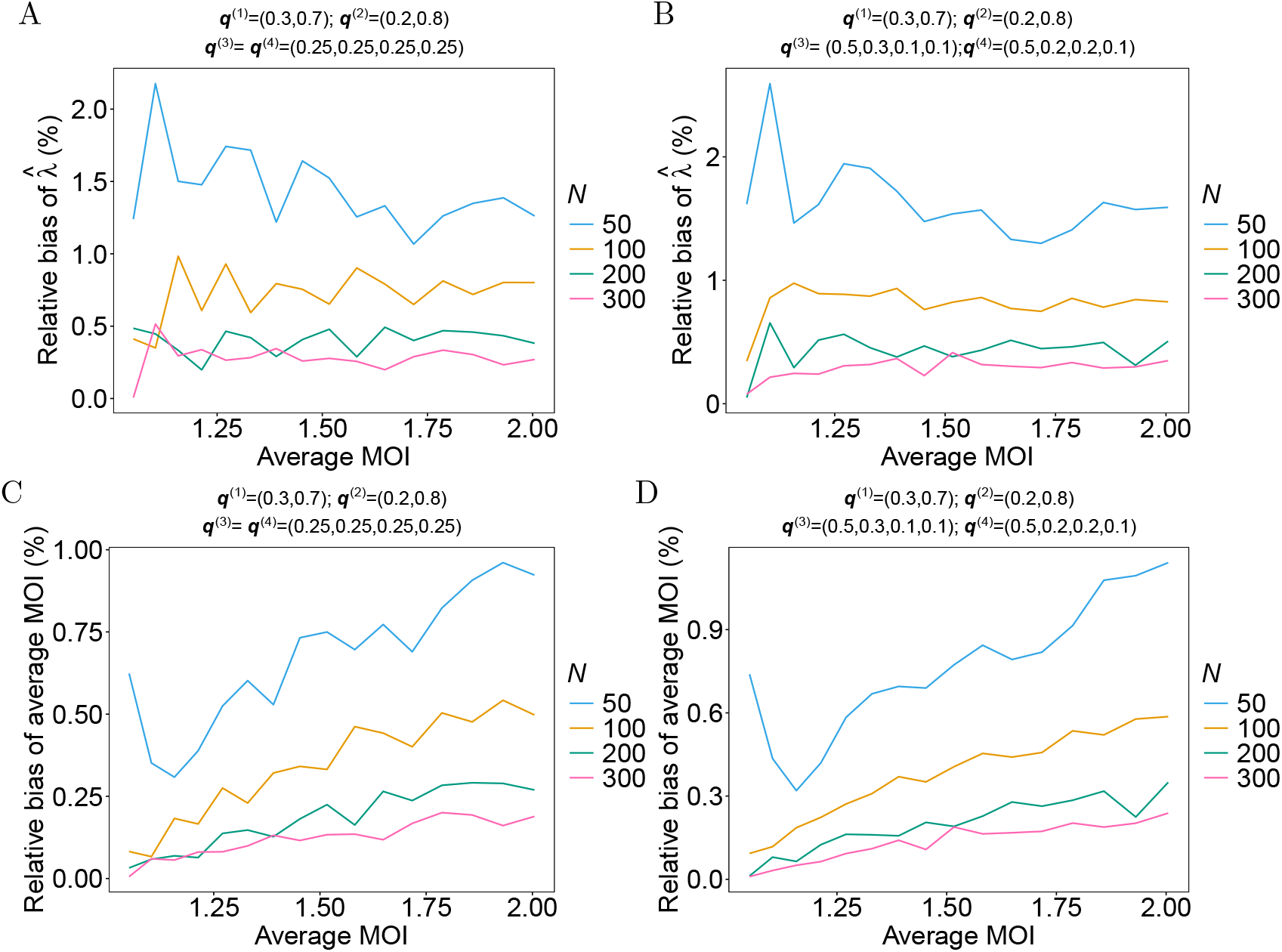
Relative bias of the estimator for MOI parameter and the average MOI as a function of the average MOI. Panels A and B show the relative bias of the estimated Poisson parameter 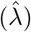, while panels C and D show the estimated average MOI 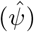 as a function of the true average MOI (*ψ*) for varying sample sizes (*N*; colors). True underlying haplotype frequencies were derived from fixed allele frequencies (shown at the top of each panel) under the assumption of linkage equilibrium. For panels A and C the neutral loci (*ℓ* = 3, 4) had balanced frequencies, while for panels B and D unbalanced frequencies were assumed.

**Fig 5.**
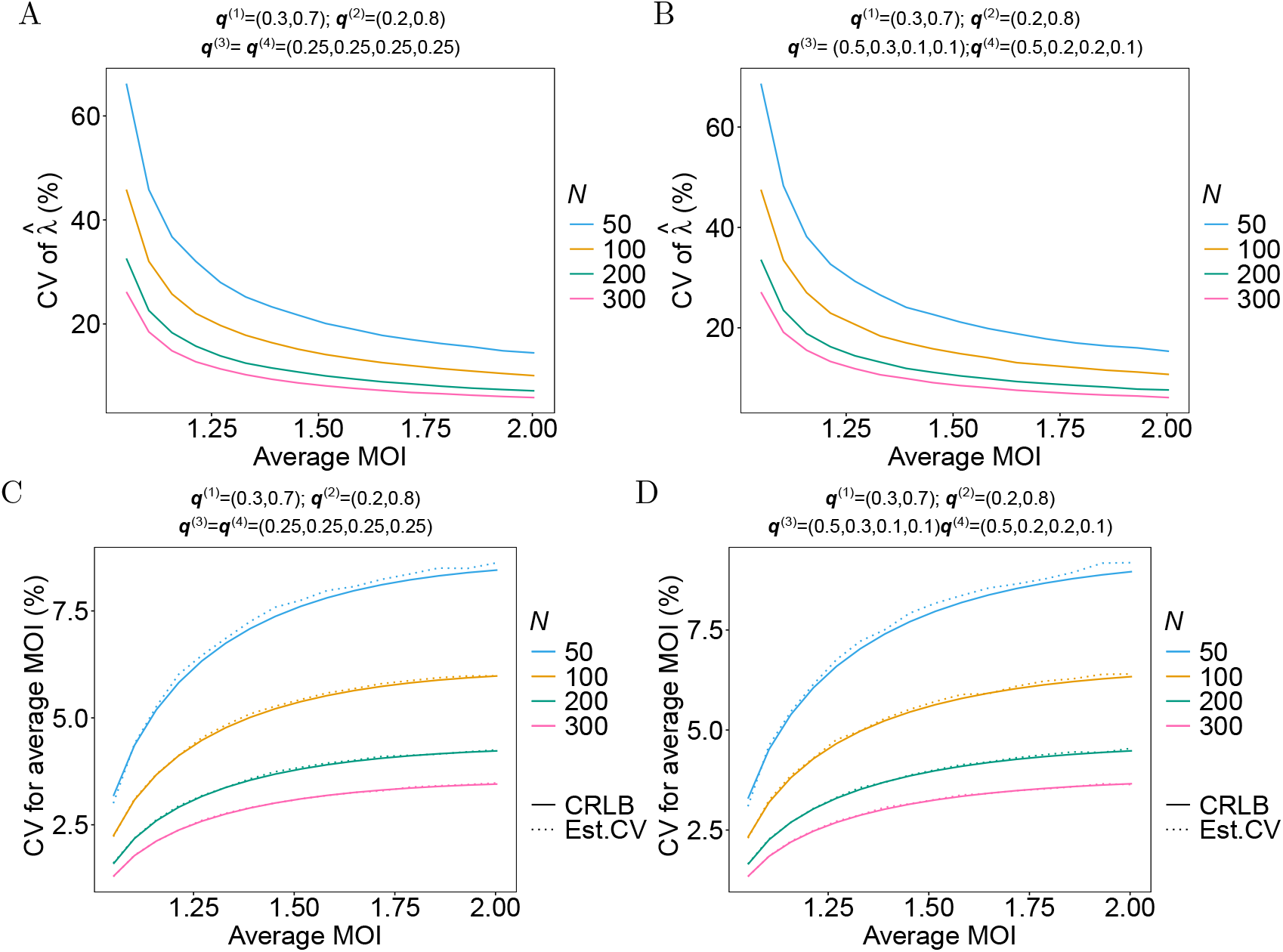
Variation in the estimated Poisson parameter 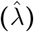 and the estimated average MOI 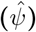 across true average MOI (*ψ*) for varying sample sizes (*N*). Different line colors represent different sample sizes. The first row shows the coefficient of variation (CV, in %) for 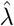 when neutral loci (*ℓ* = 3, 4) have identical (A) versus different (B) allele frequencies. The second row (C, D) compares the estimated CV (in %)(Est.CV.) of 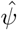 (dashed lines) to its Cramér-Rao Lower Bound (CRLB)(solid lines) under the identical (C) and different (D) allele frequency scenarios, respectively.

Here, *D* = 2 was chosen. At the two deletion markers only two ‘variants’ were assumed: the deletion and the wild type. The following notation is used: ‘1’ represents the deletion and ‘2’ the wild type. Specifically, ‘1-1’ corresponds to haplotypes with deletions at both *Pfhrp*2/3 loci, ‘1-2’ denotes haplotypes with a deletion at the first *Pfhrp*2/3 marker only, ‘2-1’ denotes haplotypes with a deletion at the second *Pfhrp*2/3 marker only, and ‘2-2’ represents haplotypes with functional variants at both loci.

Results are organized to first address estimator bias and then variability. Here we present results for the genetic architectures chosen in Simulation setup in S1 Appendix for scenarios 1 and 2. Results for the remaining scenarios are presented in S1 Appendix. As the overall trends in bias and variability are comparable across all parameters and scenarios, the additional results are reported there for completeness. Readers interested in the estimator’s performance under these additional scenarios are referred to that section.

#### Bias of haplotype frequencies

The accuracy of the marginal haplotype frequency estimates was assessed for the four haplotype as a function of different parameters.

As shown in Fig 3A,C, the magnitude of bias is generally small (typically within ±0.005) across all conditions, indicating that the estimator performs well in recovering true haplotype frequencies. As expected, larger sample sizes (*N* = 300) substantially reduced bias for all haplotypes and MOI levels. Bias increases with MOI, especially for smaller sample sizes (*N* = 50, 100), likely because the estimates are governed by outliers.

The patterns differ notably between the allele frequency configurations at the neutral markers (*ℓ* = 3, 4). In the balanced frequency scenario (Fig 3A), where neutral alleles are equally common (i.e., ***q***^(3)^ = ***q***^(4)^), the many possible haplotype combinations are all equally likely. This balance makes it easier for the model to correctly estimate the true haplotypes frequencies, resulting in very small biases. However, in the skewed frequency scenario (Fig 3C), a few specific haplotype combinations are very common, while most others are rare. Particularly, with smaller sample sizes, it becomes difficult for the model to get a clear picture of all the different haplotypes. This leads to a more noticeable bias, particularly for the most common wild types (‘2-2’).

Additionally, the consistent bias pattern observed for the wild-type haplotypes (‘2-2’) can also be attributed to the ‘masking’ of deletion haplotypes in mixed infections. When a deletion haplotype is super-infecting together with a wild-type haplotype, the functional allele from the wild-type haplotype leads to a positive genotyping result. Consequently, the presence of the deletion haplotype is masked. This causes a systematic overestimation of the wild-type haplotype frequency in the data. The model presented here is designed to correct for this bias. The results confirm its effectiveness, hence the more pronounced biases for the wild-type haplotypes. The model infers the presence of masked deletions and in the process, for smallest sample sizes (*N* = 50, 100), a slight positive and negative bias is observed, indicating that with limited data, the model lacks the power to fully correct for the ‘masking effect’. However, for larger sample sizes (*N* = 200, 300), the model effectively corrects the initial overestimation, reflecting the model’s improved ability to infer masked deletions with more data.

These results suggest that the estimator performs reliably overall, with minor biases that diminish as the sample size or MOI increases.

#### Variation of haplotype frequencies

The precision of haplotype frequency estimates was assessed using their empirical standard deviation (SD). As shown in Fig 3B,D, the SD decreases substantially for all haplotypes as sample size (*N*) increases.

The wild type (‘2-2’), being the most frequent (true frequency = 0.56), shows the highest SD across all conditions. This pattern reflects the greater absolute variability typically associated with estimating more common frequencies. In contrast, the rarer haplotypes (‘1-1’, ‘1-2’, ‘2-1’) show lower SD values, consistent with their lower true frequencies.

The effect of transmission intensity is also clear. For a given sample size, the SD generally increases as the mean MOI increases. While higher MOI provides more genetic information per host, it also introduces greater complexity in the observed data. The higher number of possible haplotype combinations in mixed infections increases the variability of the estimates. This effect is most pronounced for the most frequent haplotype (‘2-2’) and for limited data, i.e., smaller sample sizes (*N* = 50, 100).

Notably, the precision of the estimates is almost similar between the balanced and unbalanced neutral allele frequencies, particularly for sample sizes of *N* = 200, 300. This suggests that the model’s ability to precisely estimate the marginal deletion haplotype frequencies is robust to the underlying genetic diversity at neutral markers.

It is important to note that the SD values are generally small across all scenarios, typically remaining below 0.08. This indicates that the model presented provides reliable and stable estimates of haplotype frequencies, even for small sample sizes or high transmission intensity, when there is a lot of variability in the data.

#### Bias of Mean MOI

The performance in estimating the mean MOI (*ψ*) was investigated by first examining the bias of its underlying parameter, *λ*, from the conditional Poisson model. The estimate 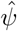 is derived from 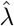 using the relationship in (35). The relative bias for both parameters is shown in Fig 4.

Fig 4A,B shows that the relative bias for 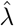 is consistently positive across all conditions, and is most pronounced for smaller sample sizes. This is because the range of the Poisson parameter has a lower, but no upper bound. Hence, rare outliers lead to overestimates, which are larger in magnitude than underestimates. Overall, the magnitude of the bias is small. It remains below 1% for sample sizes *N* ≥ 100 across all MOI values, indicating accurate estimation of the underlying Poisson parameter. Bias is similar as a function of the true average MOI for the two underlying choices of haplotype frequencies. The same results hold true if the estimator is transformed to the average MOI (Fig 4C,D). Bias is slightly larger if the allele frequencies at the neutral loci are unbalanced (cf. Fig 4A,C with B,D) irrespective of sample size.

In all cases, smaller sample sizes lead to a larger bias because the data contain less information and is more affected by sampling variability. Furthermore, the unbalanced allele frequency scenario consistently exhibits a slightly higher bias than the balanced scenario. This is because for unbalanced frequencies at neutral markers, samples are less likely to adequately reflect super-infections, leading to slight underestimates in most cases and occasional outliers yielding substantial overestimates, resulting in larger bias.

The pattern for 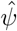 is a direct consequence of the nonlinear relationship between *λ* and *ψ*. The function *ψ*(*λ*) is increasing and convex. Therefore, the small, positive bias in 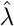 (which is almost constant for larger MOI) is amplified when transformed into 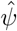. This amplification is more pronounced at higher transmission intensities where the function is steeper. Overall, these results suggest that the estimator performs very well. The observed bias in 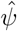 is the consequence of non-linearly transforming the bias of 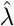. Moreover, the results indicate that the estimator is asymptotically unbiased.

#### Variation of Mean MOI

The precision of the estimates for the Poisson parameter *λ* and the mean MOI (*ψ*) was evaluated using the coefficient of variation (CV). Fig 5 shows the CV for both parameters as a function of true MOI and sample size.

A strong inverse relationship between sample size and CV is evident for both 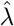 (Fig 5A,B) and 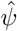 (Fig 5C,D). As expected, for both parameters, the CV decreases substantially as the sample size increases.

The effect of higher MOI is also notable. For 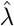, the CV generally decreases as the true mean MOI increases for a given sample size. This is an artifact of the scale of the parameter. Namely, for true values of *λ* close to 0, even a very small absolute standard deviation yields a large relative contribution (because the standard deviation is essentially divided by a value close to 0). However, the non-linear transformation into the average MOI changes this pattern. In particular, the denominator of the CV is always larger than 1. The CV of the mean MOI increases as a function of mean MOI. This occurs because higher MOI increases the probability for super-infections, and leads to more sampling variability.

Furthermore, the precision is hardly affected by the underlying haplotype-frequency distribution. The CV values are very similar when assuming balanced *vs*. unbalanced allele frequencies at the neutral markers. Although in the former case the precision is slightly higher, particularly for the estimated average MOI at smaller sample sizes.

A comparison of the CV and the (transformed) Cramér-Rao lower bound (CRLB) shows close agreement in both balanced and unbalanced neutral allele frequencies (Fig 5C,D). The differences become smaller for larger sample sizes, indicating that the estimator is efficient.

### Empirical Data Analysis

To demonstrate the practical utility of the method, it was applied to data from two studies conducted in Jagdalpur, India, refereed to as “hospital study” and “community study” (see Empirical data in Methods).

In the studies, 247 hospital patients and 104 community members were identified who had both confirmed *P. falciparum* infections and complete genetic information for all molecular markers. The method to estimate the frequency of *Pfhrp*2/3 deletions was applied to each dataset, and results marginalized to *Pfhrp*2/3 haplotypes.

Results for the hospital study are presented in Table 2. The wild-type (no deletions at any *Pfhrp*2 or *Pfhrp*3 marker) predominated at frequency 92.98% (95% CI: 91.46%–94.37%), while haplotypes carrying deletions were rare. The most common haplotype with deletions involved a single deletion in the HRP3 upstream flanking region (2.11%; 95% CI: 1.69%–2.65%). Complete deletions of all eight markers for both HRP2 and HRP3 genes occurred at frequency 1.27% (95% CI: 0.76%–1.91%). The MOI parameter was estimated as 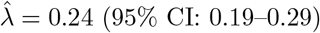, translating into an average MOI of 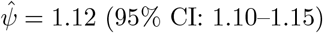, indicating moderate transmission intensity. Based on these estimates, the haplotype with complete deletions in the HRP3 region had a prevalence of 0.53% (95% CI: 0.32%–0.80%), while the complete deletion haplotype had a prevalence of 1.43% (95% CI: 0.86%–2.14%). The prevalence of infections with deletions that are not directly observable at any position (*A* = ∅, *B* = *D* = 1, …, 8 in the notation of section Infections without fully functional HRP) is 0.28% (95% CI: 0.14%–0.50%). The prevalence of infections containing any haplotype with deletions (1 − *q*_∅,∅_) is 7.82% (95% CI: 5.86%–10.17%).

**Table 2.** Estimated haplotype frequencies for hospital-based infections (*N* = 256). Shown are the eight most frequent haplotypes ranked by estimated frequency. For both *PfHRP* 2 and *PfHRP* 3, four molecular markers were analyzed (upstream flanking region, exon 1–2, exon 2, and downstream flanking region), and “WT” indicates wild type (no deletion) and “DEL” indicates deletion.

| *PfHRP2* |  |  |  | *PfHRP3* |  |  |  | Freq. % | Prev. % |
| --- | --- | --- | --- | --- | --- | --- | --- | --- | --- |
| up | ex 1-2 | ex 2 | down | up | ex 1-2 | ex 2 | down | (95% CI) | (95% CI) |
| WT | WT | WT | WT | WT | WT | WT | WT | 92.98<br>(91.46–94.37) | 93.73<br>(92.07–95.14) |
| WT | WT | WT | WT | DEL | WT | WT | WT | 2.11<br>(1.69–2.65) | 2.37<br>(1.90–2.97) |
| DEL | DEL | DEL | DEL | DEL | DEL | DEL | DEL | 1.27<br>(0.76–1.91) | 1.43<br>(0.86–2.14) |
| DEL | WT | WT | WT | DEL | WT | WT | WT | 0.93<br>(0.56–1.40) | 1.04<br>(0.63–1.57) |
| WT | WT | WT | WT | DEL | DEL | DEL | WT | 0.92<br>(0.55–1.38) | 1.03<br>(0.62–1.55) |
| DEL | WT | WT | WT | WT | WT | WT | WT | 0.89<br>(0.53–1.34) | 1.00<br>(0.60–1.50) |
| WT | WT | WT | WT | DEL | DEL | DEL | DEL | 0.47<br>(0.28–0.71) | 0.53<br>(0.32–0.80) |
| DEL | DEL | DEL | DEL | WT | WT | WT | WT | 0.43<br>(0.26–0.65) | 0.48<br>(0.29–0.73) |

In the community study, deletions were found to be more common (cf. Table 3). The wild-type haplotype had a frequency of 72.18% (95% CI: 68.57%–75.50%). The most frequent haplotypes with deletions (6.01%;95% CI: 4.21%–8.14%) had deletions in the upstream regions of both HRP2 and HRP3. Complete deletions at all eight markers for both genes occurred at 5.33% (95% CI: 3.73%–7.15%) frequency. The estimated MOI parameter was slightly lower in the community setting compared to the hospital at 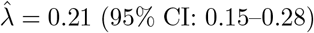, with the corresponding average MOI of 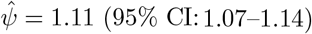. Based on these estimates, the haplotype with a single deletion in the HRP3 upstream flanking region had a prevalence of 4.53% (95% CI: 3.17%–6.13%), while the complete deletion haplotype had a prevalence of 5.88% (95% CI: 4.12%–7.93%). The prevalence of infections with deletions that are not directly observable at any position (*A* = ∅, *B* = *D* = 1, …, 8) is 0.84% (95% CI: 0.42%–1.31%). The prevalence of infections containing any haplotype with deletions (1 − *q*_∅,∅_) is 30% (95% CI: 25.5%–34.8%).

**Table 3.** Estimated haplotype frequencies for community based data (*N* = 104). Same as in Table 2, but for the community based data.

| *PfHRP2* |  |  |  | *PfHRP3* |  |  |  | Freq. % | Prev. % |
| --- | --- | --- | --- | --- | --- | --- | --- | --- | --- |
| up | ex 1-2 | ex 2 | down | up | ex 1-2 | ex 2 | down | (95% CI) | (95% CI) |
| WT | WT | WT | WT | WT | WT | WT | WT | 72.18<br>(68.57–75.50) | 74.30<br>(70.80–77.78) |
| DEL | WT | WT | WT | DEL | WT | WT | WT | 6.01<br>(4.21–8.14) | 6.64<br>(4.65–9.00) |
| DEL | DEL | DEL | DEL | DEL | DEL | DEL | DEL | 5.33<br>(3.73–7.15) | 5.88<br>(4.12–7.93) |
| WT | WT | WT | WT | DEL | WT | WT | WT | 4.10<br>(2.93–5.22) | 4.53<br>(3.17–6.13) |
| DEL | DEL | DEL | WT | WT | WT | WT | WT | 2.14<br>(1.28–3.21) | 2.38<br>(1.42–3.56) |
| DEL | WT | WT | WT | DEL | DEL | DEL | WT | 2.08<br>(1.25–3.10) | 2.31<br>(1.39–3.44) |
| DEL | DEL | DEL | WT | DEL | DEL | DEL | WT | 1.96<br>(0.98–3.14) | 2.18<br>(1.09–3.49) |
| WT | WT | WT | WT | DEL | DEL | DEL | WT | 1.12<br>(0.67–1.60) | 1.24<br>(0.74–1.82) |
| DEL | DEL | DEL | DEL | WT | WT | WT | WT | 1.06<br>(0.53–1.70) | 1.18<br>(0.59–1.88) |
| DEL | DEL | DEL | WT | DEL | WT | WT | WT | 1.03<br>(0.52–1.65) | 1.15<br>(0.60–1.82) |
| DEL | WT | WT | WT | WT | WT | WT | WT | 1.02<br>(0.51–1.63) | 1.13<br>(0.60–1.78) |
| DEL | DEL | WT | WT | DEL | WT | WT | WT | 1.02<br>(0.62–1.67) | 1.13<br>(0.56–1.81) |
| DEL | DEL | DEL | DEL | DEL | DEL | DEL | WT | 0.94<br>(0.57–1.49) | 1.05<br>(0.62–1.68) |

## Discussion

Molecular surveillance has become a cornerstone of state-of-the-art malaria control, providing critical insights into parasite diversity, transmission dynamics, and the spread of clinically significant mutations [34, 35]. The WHO lists the emergence and spread of *P. falciparum* parasites carrying *Pfhrp*2/3 deletions as one of four key biological threats in malaria control [1]. *Pfhrp*2/3 encode for the antigen targets of the most widely used RDTs. The WHO recommends to change RDT policies in areas, where the prevalence of deletions in symptomatic infections exceed 5%. This threshold-based policy highlights the requirement to accurately estimate the frequency and prevalence of *Pfhrp*2/3 deletions [21, 36].

To mitigate this gap, we introduced a novel statistical method to estimate the frequencies and prevalence of *Pfhrp*2/3 deletions, accounting for the possibilities that deletions might not be directly observable in the case of polyclonal infections. The model presented here explicitly incorporates haplotypes with deletions at one or several positions in the *Pfhrp*2/3 genes. The statistical method is based on molecular data, which contains information from markers within the *Pfhrp*2/3 genes (these can be deleted) and neutral molecular markers at which no deletions occur. This allows probabilistically reconstructing whether deletions are unobservable within infections. The statistical model is exact in the sense that it incorporates all possible haplotypes that are potentially circulating in the parasite population based on the observations. The method is based on the exact likelihood function, i.e., without numerical simplifications. In this context, MOI refers to the number of super-infections (i.e., one or more independent infectious events, with one pathogen haplotype being transmitted at each of them) and ignores co-infections (i.e., the co-transmission of multiple pathogen variants at the same infective event). While super-infections approximate co-infections, the advantage is that mosquito population (or rather the distribution of mosquitoes carrying specific sets of pathogen variants) does not need to be modeled explicitly. The latter would require several additional modeling assumptions. The distribution of MOI is flexible in the model formulation. However, for maximum-likelihood estimation, specific assumptions regarding the MOI distribution are necessary, as the set of model parameters needs to be well defined.

Here, following [] MOI was assumed to follow a conditional Poisson distribution. It is acknowledged that alternative distributions could be considered to better reflect heterogeneity in exposure [37]. For example, the negative binomial distribution, which incorporates a dispersion parameter, can accommodate over-dispersed MOI data resulting from heterogeneous biting rates across hosts. However, this approach introduces the challenge of reliably estimating both the mean and dispersion parameters simultaneously, as argued in [38]. As an alternative, a mixed Poisson distribution could be applied by assigning different infection rates to population strata, though this requires accurate stratification of data that may not be readily available in field settings. A more flexible, non-parametric approach to estimating the MOI distribution directly from the data without assuming a specific parametric form has also been proposed in [38]. Nevertheless, it has been demonstrated that the conditional Poisson framework used here is sufficient to capture the essential features of MOI [38].

Because of the complexity of the likelihood function, it was not possible to derive an explicit solution for the estimator. Consequently, the EM-algorithm was adapted to numerically obtain the MLE. An advantage of the statistical model is its ability to accommodate complex genetic architectures. Specifically, the method is not restricted to a particular type of molecular data or assumes bi-allelic markers.

It was shown by simulation that the estimator possesses desirable finite-sample and asymptotic properties. The estimates for haplotype frequencies and the average multiplicity of infection (MOI) showed little bias across a range of model parameters, with accuracy and precision improving when increasing sample size. In fact, the results suggest that the estimator is asymptotically unbiased. Notably, the estimator performed reliably even for unbalanced marginal allele frequency distributions at neutral markers, which occurs commonly in natural parasite populations [39, 40]. The precision of the estimator was high (quantified by their standard deviation and coefficient of variation), and aligned with theoretical expectations. More precisely, the empirical variance of the mean MOI estimator was found to coincide with the Cramér–Rao lower bound, confirming its statistical efficiency and indicating that maximum possible information is extracted from the data [31, 32].

The primary purpose of this statistical method is the estimation of haplotype frequencies with *Pfhrp*2/3 deletions. These frequencies (relative abundance in parasite population) are intrinsically linked to their prevalence (probability of occurring in an infection) via MOI. Several formulae for prevalence were derived to assess how likely infections are in which deletions can be directly observed, and how common deletions cannot be directly observed. Besides the functional relationship between frequency and prevalence, the estimation of MOI represents a particularly important epidemiological metric. As a direct correlate of transmission intensity, MOI serves as a robust measure for evaluating the impact of malaria control interventions [24, 41]. Unlike routine case counts, which can be influenced by variations in healthcare seeking behavior (or access) and diagnostic practices, MOI provides a well-defined metric. A sustained reduction in MOI over time can therefore offer early evidence of successful transmission reduction, even before corresponding decline trends are visible in incidence data [42].

The successful application to empirical data from a tribal community in India illustrates the practical utility of the model. It yielded stable and interpretable estimates of both haplotype frequencies and transmission intensity.

Despite its strengths, the method has limitations. The most significant constraint is computational, arising from the need to recreate all possible haplotypes that could generate the observed genetic data from a mixed infection. The number of these combinations grows exponentially with the number of loci and alleles observed per marker, which could pose challenges for highly polyclonal infections with many polymorphic markers. This could pose a computational constraint for a large number of polymorphic markers.

Additionally, in the current framework, all missing alleles at the *Pfhrp*2/3 markers are attributed to true genetic deletions. In reality, allele absence could also result from low parasite density, whereby the parasite might be present in the blood at levels too low to be detected by the assay (especially early in an infection or in the case of a low density infection), assay failure or human error [36, 43]. A future extension could jointly model the probability of a genuine deletion and the probability of technical assay failure, thereby enhancing robustness in settings with variable laboratory conditions. However, this would require additional validation data, such as confirmatory sequencing to distinguish between deletion and failure.

In any case, the method presented here enables malaria control programs to generate reliable evidence based estimates of *Pfhrp*2/3 deletion prevalence, which are essential for informing diagnostic policy decisions. It also provides a robust, scalable tool for monitoring transmission intensity through MOI estimation, supporting the evaluation and refinement of malaria control strategies in diverse epidemiological settings.

## Methods

Here, a statistical model to accurately estimate the population frequency distribution of *Pfhrp*2 and/or *Pfhrp*3 deletions in the presence of mixed clone infections is introduced. More precisely, the statistical framework used molecular data at markers in the *Pfhrp*2 and/or *Pfhrp*3 genes and molecular information from neutral markers to derive the frequency distribution of haplotypes circulating in the population as well as a population level estimate of multiplicity of infection (MOI). Following [23, 24] MOI is formally defined as the number of super-infections, i.e., independent malaria infections during one disease episode.

First, a mathematical description of the statistical model is presented, including the formal definition of MOI, haplotypes, their genetic architecture, and structure of molecular data.

Point estimates for the MOI and haplotype distribution are obtained from the statistical model via maximum-likelihood. Since the model does not allow for an explicit solution, the expectation maximization algorithm is used to derive the maximum likelihood estimate (MLE) of the model parameters.

### Model setup

A genetic architecture of *L* multi-allelic markers (loci) is considered. The *n*_*ℓ*_ alleles segregating at marker *ℓ* are denoted by 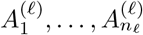. The first *D* markers correspond to markers, e.g., exons, in *Pfhrp*2/3. The convention is used that the first allele 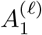 at loci 1, …, *D* corresponds to a deletion, while the remaining alleles are distinct variants.

With this genetic architecture in mind, a parasite haplotype ***h*** is characterized by specific allelic variants at the *L* markers. Formally, a haplotype is denoted as a vector ***h*** = (*h*_1_, …, *h*_*L*_), where *h*_*ℓ*_ ∈ {1, …, *n*_*ℓ*_}.. The set of all 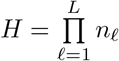 possible haplotypes is denoted by 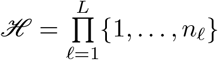

A haplotype ***h*** with deletions at all of the first *D* markers is referred to as “complete deletion”. A haplotype with deletions at some but not all of the first *D* markers is referred to as “partial deletion”. The advantage of partitioning the genetic architecture at *Pfhrp*2/3 into *D* markers is that partial deletions can be easily modeled (cf. Fig 1).

Sometimes, it is more convenient to label haplotypes by numbers 1, …, *H*, rather than using their vector representation. Hence, each haplotype ***h*** is identified with its mixed-radix representation, i.e., haplotype ***h*** is identified with its label

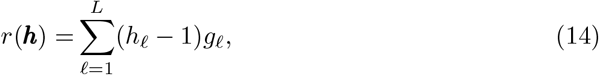

where

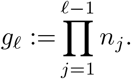

The sporozoite stage in the mosquito salivary glands serves as the census point for the haplotype frequency distribution [22], which are denoted by the vector ***p*** := (*p*_***h***_)_***h***∈ℋ_ = (*p*_1_, …, *p*_*H*_), where *p*_***h***_ = *p*_*h*_ (if *h* = *r*(***h***)) is the frequency of haplotype ***h***.

Parasites are observed in human hosts during the erythrocytic cycle. A confounding factor is that hosts can be super-infected due to multiple disease exposures. Here, super-infections refer to independent infective events during one disease episode. It is further assumed that at each infecting event, exactly one parasite haplotype is transmitted to the host. This ignores the possibility of co-infections, i.e., the transmission of multiple parasite haplotypes during one infective event (for more discussion on this assumption, see e.g. [29]). Following [22, 24, 30] the number of super-infections is referred to as multiplicity of infection (MOI).

Assuming infections are rare and independent, the number of infections per individual, denoted by *m*, is a realization of a random variable following a Poisson distribution. Considering only disease-positive hosts, MOI follows a conditional or positive Poisson distribution (*m* ~ CPoiss(*λ*)), namely

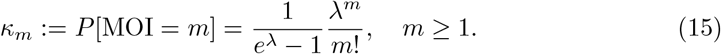

The mean MOI, denoted by *ψ*, is calculated to be

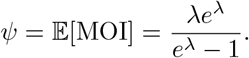

Notably, the derivation of the statistical model and the likelihood function in the following are independent of the assumption of the Poisson distribution and can be changed by any other distribution of the positive integers. However, derivation of the EM algorithm, is based on the Poisson assumption.

The assumption of exactly one parasite haplotype being transmitted at each infectious event, implies that, given MOI *m*, the configuration of haplotypes infecting a host follows a multinomial distribution with parameters *m* and ***p***. Given MOI = *m*, let *m*_***h***_ denote the number of times the host was infected with haplotype ***h*** (subject to the constraint 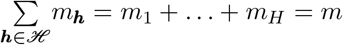). Thus, an infection is characterized by a vector ***m*** := (*m*_***h***_)_***h***∈ℋ_ with |***m***| := *m*_1_ + … + *m*_*H*_ = *m* follows a multinomial distribution (***m*** ~ Multi(*m*, ***p***)), i.e.,

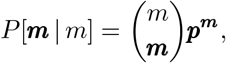

where 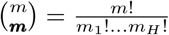 and 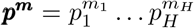.

In practice, MOI is unobservable and haplotype information is typically unavailable because molecular assays yield unphased information (cf. Fig 2). More precisely, at each marker *ℓ* the set of alleles *x*_*ℓ*_ ⊆ {1, …, *n*_*ℓ*_*}* present in the infection is observable, it is in general unknown which haplotypes are present in an infection (i.e., for which haplotypes ***h*** the number *m*_***h***_ *>* 0). As the first allele 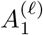 corresponds to a deletion at markers *ℓ* = 1, …, *D*, which is by nature “unobservable”, the convention 1 ∈ *x*_*ℓ*_ for *ℓ* = 1, …, *D* is made here. Formally, in a clinical sample at marker *ℓ* the observable information is

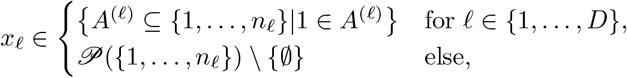

with 𝒫 denoting the power set. (Note, the empty set is excluded because only disease positive samples are considered.) The set of all possible observations is

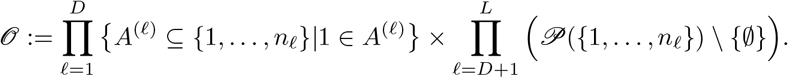

An observation ***y*** a called a sub-observation of ***x***, denoted ***y*** ⪯ ***x***, if all alleles observed in ***y*** are also observed in ***x***. Formally,

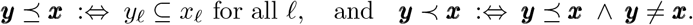

The relation “⪯” defines a partial order on 𝒪. The set of all sub-observations of ***x*** is defined by

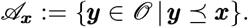

Remember, in general, it is unknown which haplotypes are present in an infection, since only the observation ***x*** is known. From an observation ***x*** the presence of certain haplotypes in the infection can be ruled out, as these are incompatible with the observation ***x***. The set of all haplotypes, “compatible” with observation ***x***, is denoted by

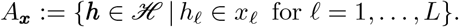

Note, if a haplotype ***h*** is an element of *A*_***x***_ it does not imply that haplotype ***h*** was present in the infection leading to observation ***x***, rather its presence cannot be ruled out. Moreover, for a sub-observation ***y*** ⪯ ***x***, i.e., ***y*** ∈ 𝒜_***x***_, any haplotype ***h*** compatible with ***y*** is also compatible with ***x***, i.e., if ***y*** ∈ 𝒜_***x***_, then ***h*** ∈ *A*_***y***_ ⇒ ***h*** ∈ *A*_***x***_.

Note that for a given MOI *m* and observation ***x***, there may exist multiple MOI vectors ***m*** that can generate the observation ***x***. Let the haplotype-to-observation mapping be defined as

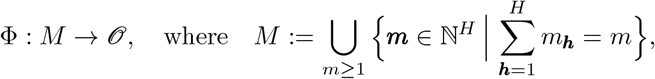

such that Φ(***m***) = ***x*** maps an MOI vector ***m*** to its observable allele set ***x***. Then the set of valid MOI vectors for observation ***x*** is defined as

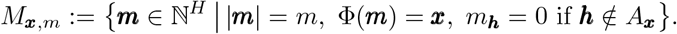

Importantly, the sets *M*_*x,m*_ are pairwise disjoint, and their disjoint union is denoted by

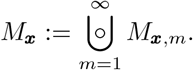

Assuming an infection with MOI *m*, the probability of observing ***x*** is

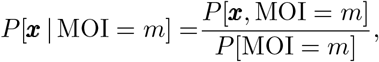

where *P*[*x*, MOI = *m*] is the probability of observation ***x*** in an infection with MOI = *m*. Therefore, using definition (15),

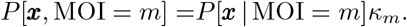

The law of total probability yields

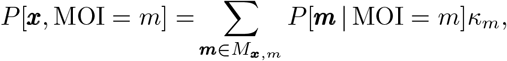

and

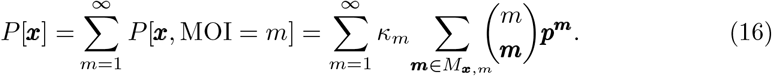

For simplicity, *P*[***x***] is denoted by *P*_***x***_ in the following. Note that ***y*** ⪯ ***x*** is equivalent to ***y*** ∈ 𝒜_***x***_. Therefore, the set *M*_*x,m*_ can be rewritten as

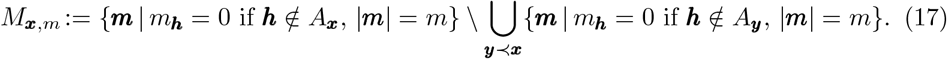

Here, {***m***|*m*_***h***_ = 0 if ***h*** ∉ *A*_***x***_, |***m***| = *m}* is the set of all possible infections with haplotypes which are compatible with ***x***, i.e., they lead to either observation ***x*** or are a proper sub-observation ***y*** ≺ ***x***. Furthermore, 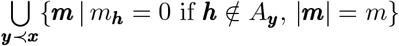 is the set of all possible haplotypes compatible with ***x***, which lead to a proper sub-observation of ***x***. The inclusion-exclusion principle yields

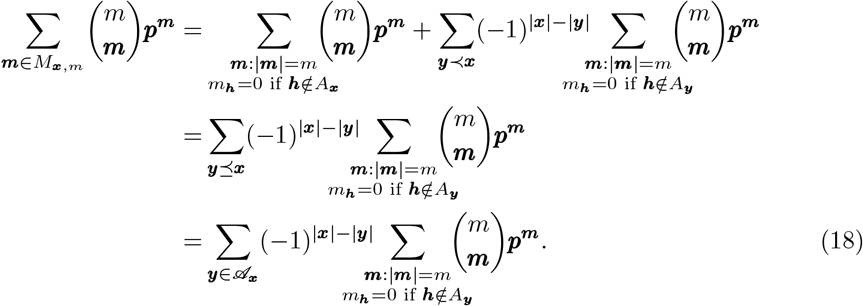

Therefore, the probability of observing ***x*** in (16) becomes

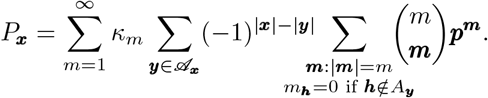

An application of the multinomial theorem and an exchange of sums give

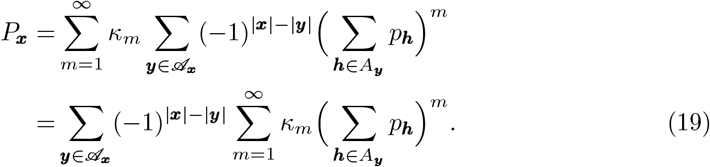

Since

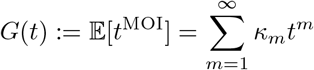

is the Probability Generating Function (PGF) of the MOI, the probability of observing ***x*** becomes,

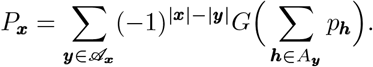

When assuming that MOI follows a conditional Poisson distribution, then the PGF is given by

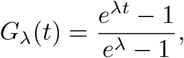

with the subscript *λ* indicating that the PGF is characterized by the Poisson parameter. In the case of the Poisson model, then

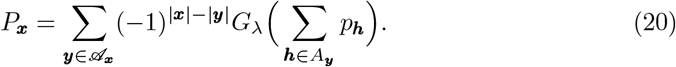

In this case, the parameter space of the model is a *H*-dimensional space Θ defined as

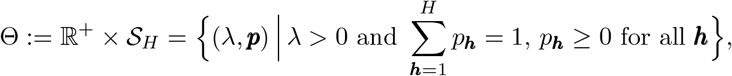

where 𝒮_*H*_ denotes the *H* − 1-dimensional simplex.

### Datasets and likelihood function

Assume a dataset 𝒳 consisting of *N* observations. The *j*-th observation is denoted by 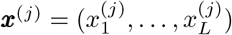, where 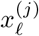 is the allelic information at marker *ℓ* in sample *j*. The number of times observation ***x*** occurs in the dataset 𝒳 is denoted by *n*_***x***_, such that

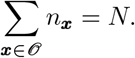

Given 𝒳, the model parameters ***θ*** := (*λ*, ***p***) ∈ Θ can be collectively estimated by the maximum-likelihood method. Since the *N* observations are assumed to be independent, the likelihood function ℒ_𝒳_ (***θ***) is

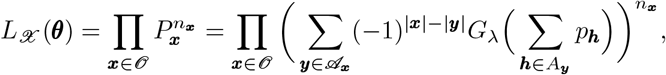

and the resulting log-likelihood function is

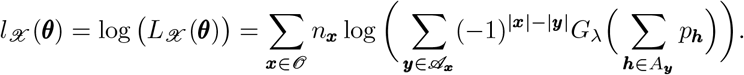

The complexity of the log-likelihood function does not allow for a closed-form solution of the maximum-likelihood estimator. Hence, for a given dataset 𝒳, the maximum-likelihood estimate (MLE) needs to be numerically derived. Here, the Expectation-Maximization algorithm (EM algorithm) is used to derive the MLE (see section Maximizing the likelihood function with the Expectation-Maximization (EM) algorithm). In general, it is a stable and fast-convergent algorithm to find MLEs [27, 44].

### Maximizing the likelihood function with the Expectation-Maximization (EM) algorithm

In this section, the EM algorithm for numerical computation of the maximum likelihood estimates (MLEs) for the model parameters ***θ*** := (*λ*, ***p***) ∈ Θ is derived. The EM algorithm is a two-step iterative method that guarantees to find a maximum of the likelihood function by alternating between an expectation step (E-step) and a maximization step (M-step) [44].

#### Expectation (E)-step

Each observation ***x*** in a dataset 𝒳 originates from an unobservable infection represented by ***m***. Hence, the observed information are ***x***^(1)^, …, ***x***^(*N*)^, collectively denoted as 𝒳, and the corresponding to unobserved information ***m***^(1)^, …, ***m***^(*N*)^ is jointly denoted by ℳ.

The expectation of the log-likelihood function of the parameter vector ***θ*** given the observable (𝒳) and unobservable (ℳ) information taken with respect to the conditional distribution of the unobservable information ℳ given the observable information 𝒳 and parameter vector ***θ***^(*t*)^ is called the *Q*-function. Formally, it is defined by

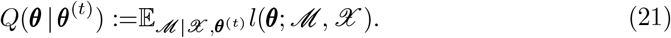

Here, the parameter vector ***θ*** is considered a variable, while the choice ***θ***^(*t*)^ is regarded to be constant. Because observations are independent, the above simplifies to

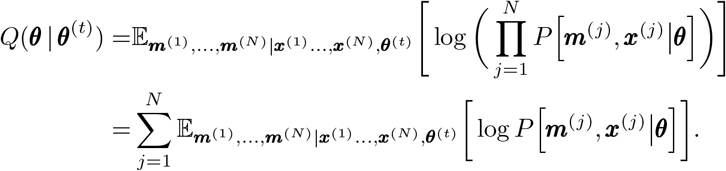

Straightforward marginalization (cf. [25] for a detailed derivation) yields

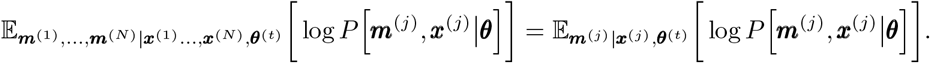

Therefore, the *Q*-function becomes

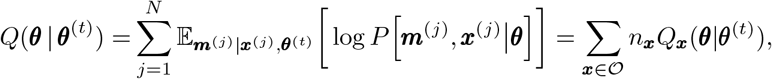

where

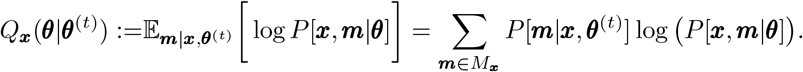

This yields

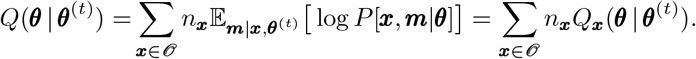

Using the definitions of conditional probabilities one obtains

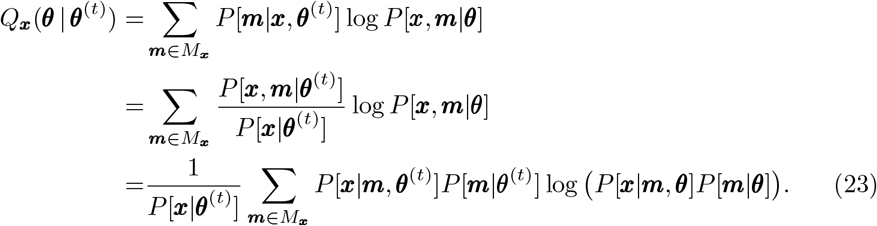

Since the sum in (23) runs over all ***m*** ∈ *M*_***x***_ and any such vector ***m*** yields observation ***x***, the identity *P*[***x***|*m*, ***θ***^(*t*)^] = 1 holds. Moreover, by the Theorem of Total Probability

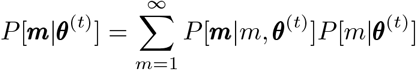

so that, by recalling the definition of *M*_*x,m*_ (17),

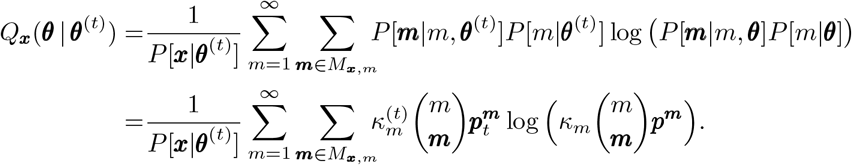

The inclusion-exclusion principle from (18) yields

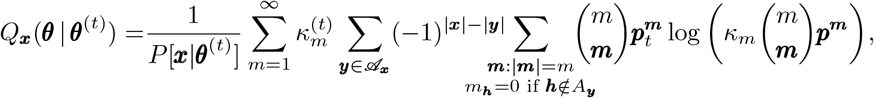

and

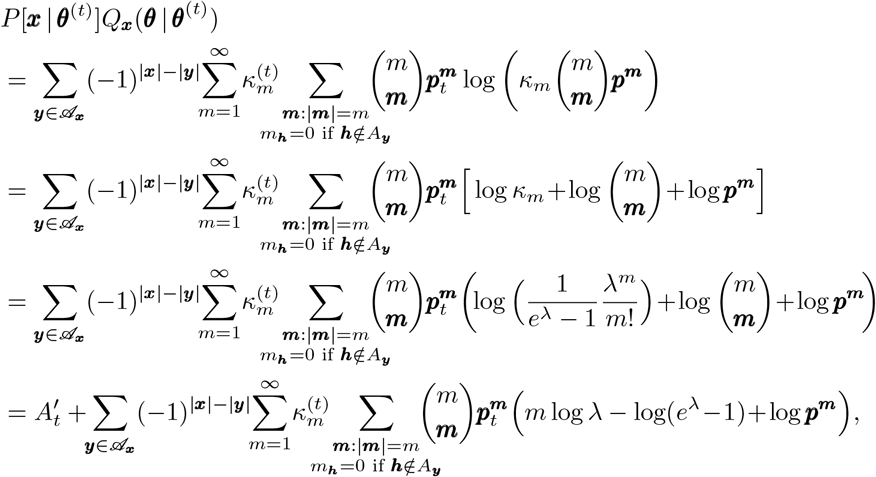

where

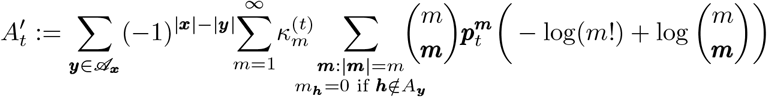

is independent of the parameters ***θ***. Therefore,

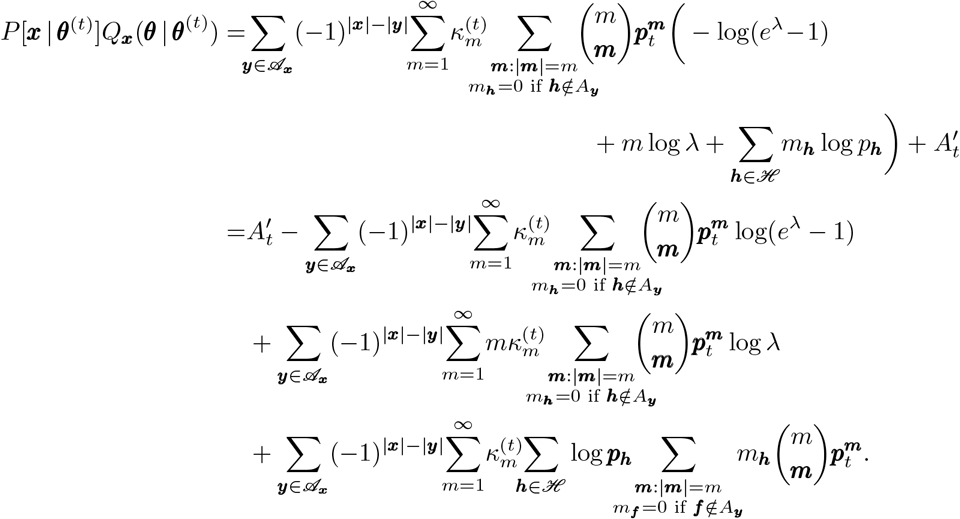

To simplify the last row above, let ***m***_−***h***_ denote the MOI vector ***m*** with the component of haplotype ***h*** reduced by 1, i.e., *m*_***h***_ is replaced by *m*_***h***_ − 1, which allows rearrangement of the binomial coefficient. Furthermore, using the definition the expression of *P*[***x*** | ***θ***^(*t*)^] from (19) gives

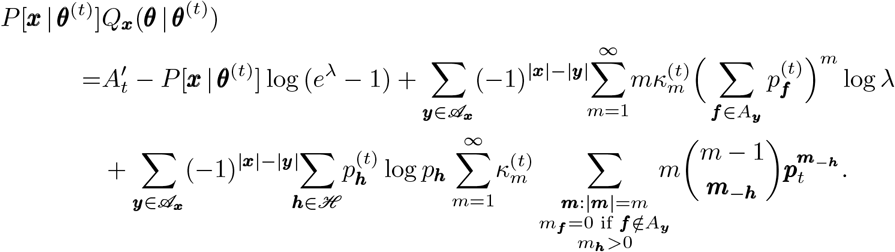

Taking one factor in the third line outside the second sum and simplification of the third line gives

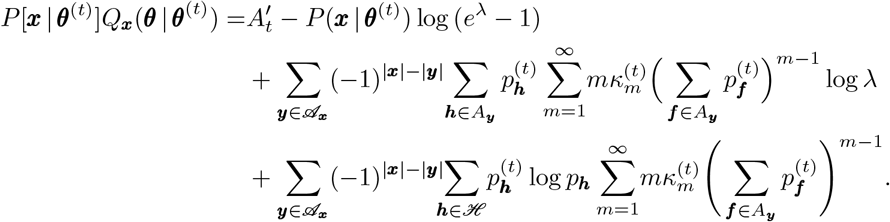

The first derivatives of the PGF occur in the above expression. Namely, for ***h*** ∈ *A*_***y***_ the derivative of the PGF gives

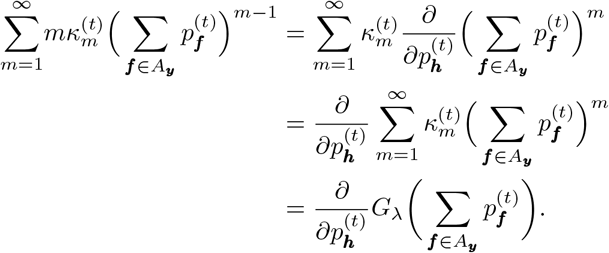

Hence,

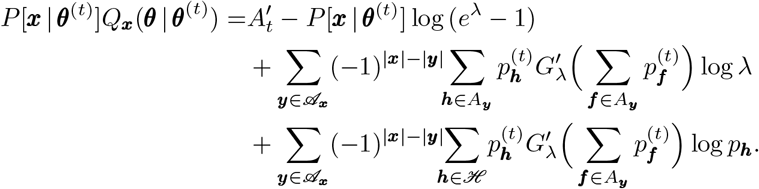

The factor log *p*_***h***_ occurs in the last row. Since ultimately the *Q*-function is maximized with respect to *p*_***h***_, it is convenient to introduce the indicator function of the set *A*_***y***_

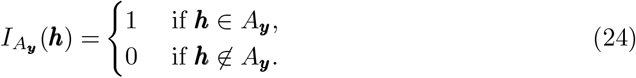

Hence,

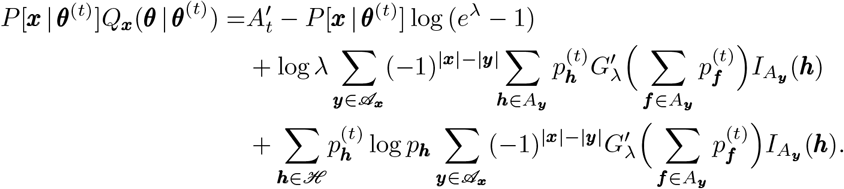

Therefore,

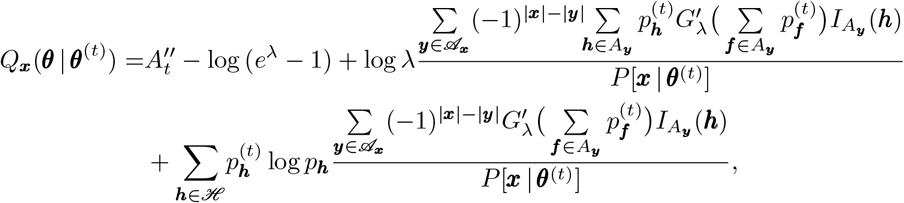

With 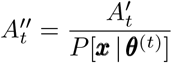. The definition of *P*[***x*** | ***θ***^(*t*)^] in (20) yields,

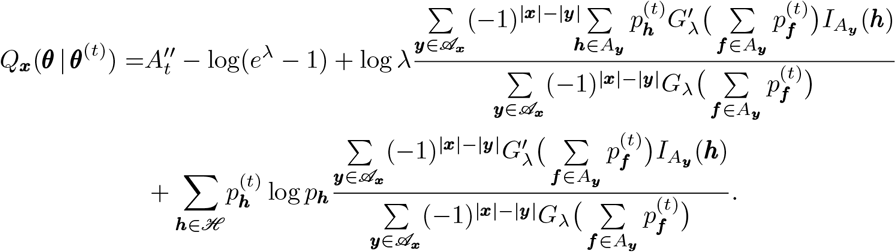

The actual *Q*-function in (23) becomes

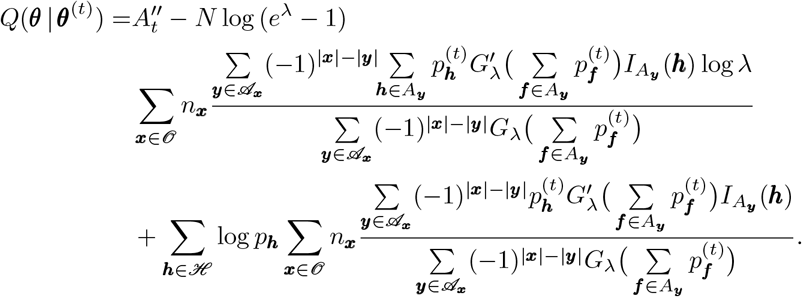

By defining

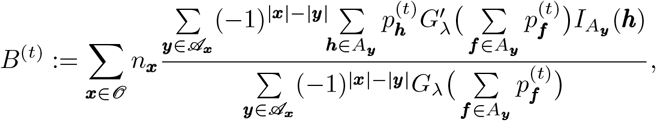

and

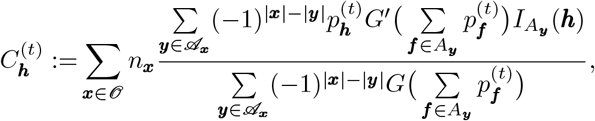

the *Q*-function is rewritten as

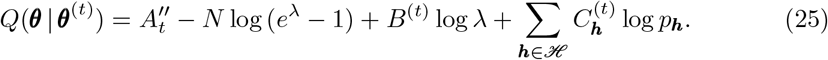

This *Q*-function is maximized in the “Maximization step” of the EM algorithm.

#### Maximization (M)-step

In the M-step of the EM-algorithm, the *Q*-function (25) is maximized with respect to the parameters ***θ***. This leads to the updated parameter estimates

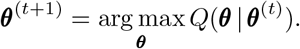

Because the haplotype frequencies are an element of the (*H* − 1)-dimensional simplex, it is convenient to maximize the *Q*-function using the method of Lagrange-multipliers [45] under the constraint 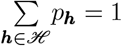. The parameter space for this constrained optimization is extended to include a Lagrange multiplier *β*. Define

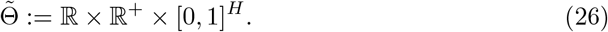

Let the extended parameter vector be defined as

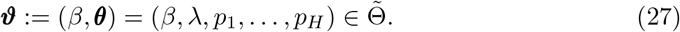

The Lagrangian function to be maximized is then

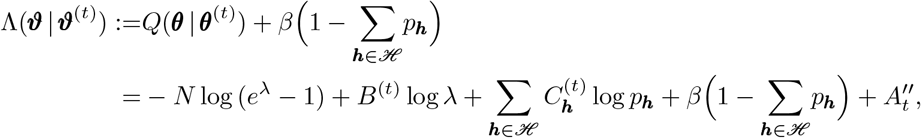

where *β* is the Lagrange multiplier. The parameters update ***θ***^(*t*+1)^ is obtained by omitting the Lagrange multiplier from ***ϑ***^(*t*+1)^, where the latter is the solution of

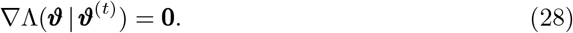

The components of ∇Λ(***ϑ*** | ***ϑ***^(*t*)^) are

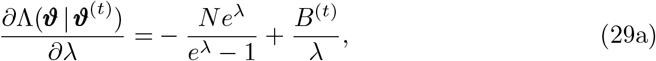

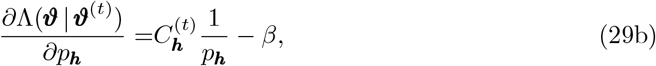

and

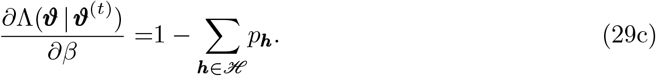

Equating (29b) to zero gives

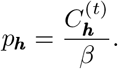

Substituting this expression into (29c) implies

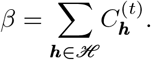

Thus the updated frequency of haplotype ***h*** in step *t* + 1 of the EM-algorithm is

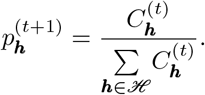

From (29a) the updated MOI parameter *λ* is obtained by solving

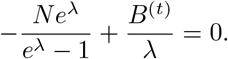

This non-linear equation has no closed-form solution but can be numerically solved using a 1-dimensional Newton-Raphson method [46]. Let

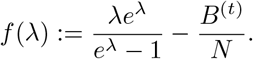

The 1-dimensional Newton-Raphson method yields the recursive equation

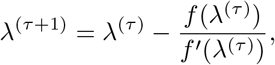

where

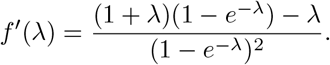

Therefore, the recursive equation is given by

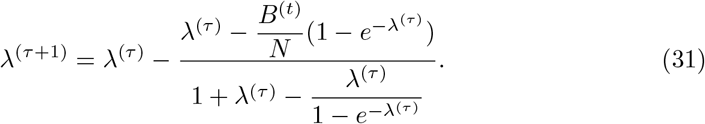

The updated MOI parameter *λ*_*t*+1_ is the value *λ*^(*τ*)^ for which convergence of (31) is reached numerically.

#### The algorithm

The EM algorithm leads to the following iteration. First, an arbitrary initial condition, *λ*^(0)^, ***p***^(0)^, is chosen. In the first step of iteration *t* + 1, the haplotype frequencies are updated as

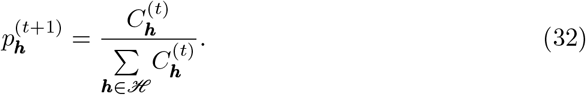

In the second step, the MOI parameter is updated by iterating

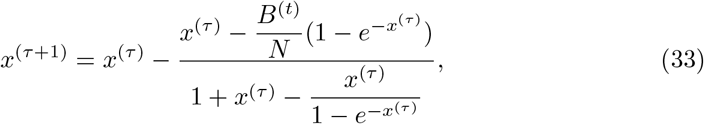

from the initial condition *x*^(0)^ = *λ*^(*t*)^ until convergence is reached numerically, i.e., until the condition |*x*^(*τ*+1)^ − *x*^(*τ*)^ |*< ε* holds. The MOI parameter is then updated as *λ*^(*t*+1)^ = *x*^(*τ*+1)^. The above two steps are repeated until numerical convergence, i.e., until the condition 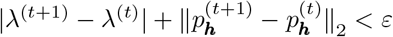 is satisfied. Upon convergence, the MLEs for the parameters are then given by

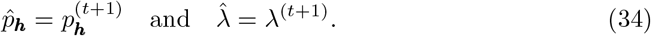

The resulting estimates provide the final parameter estimates in the EM algorithm.

### Assessing finite-sample properties by numerical simulations

The performance of the estimators under varying sample sizes and parameter choices was ascertained using Monte Carlo simulations. The goal was to assess the accuracy and precision of the estimators for the haplotype frequencies ***p*** and the average MOI, *ψ*, across a range of parameter choices.

#### Parameter choices and dataset generation

Regarding parameter choices, first a genetic architecture was specified. More precisely, the number of markers *L*, the number of markers with *Pfhrp*2/3 deletions *D* as well as the number of alleles segregating at each marker were chosen. Next, the set of circulating haplotypes together with their corresponding frequency distribution ***p*** were specified. The choices for the genetic architectures are presented in detail in section Simulation setup in S1 Appendix. For each frequency distribution ***p*** and corresponding genetic architecture, the Poisson parameter *λ* was varied across the range *λ* = (0.1, …, 1.6), corresponding to approximate average MOI values of *ψ* = (1.05, …, 2). For each set of parameters, datasets of sample size *N* = 50, 100, 200, 300 were generated.

For a parameter choice ***θ*** = (*λ*, ***p***), a dataset 𝒳 of size *N* (***x***^(1)^, …, ***x***^(*N*)^) was generated as follows. For the *j*-th observation ***x***^(*j*)^, the number of haplotypes *m* super-infecting (MOI) was drawn from a conditional Poisson distribution with parameter *λ*. Given this MOI, *m* haplotypes were sampled from a multinomial distribution with probability distribution with parameters *m* and ***p***. The haplotypes ***h*** with *m*_***h***_ *>* 0 were present in the infection. From this set of haplotypes, the alleles present at each marker were retained. This yielded the observation 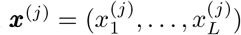, accounting for the fact that deletions are not observable, i.e., enforcing 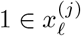 for *ℓ* = 1, …, *D*. For each set of parameters *S* = 20 000 such datasets 𝒳^(1)^, …, 𝒳^(*S*)^ were generated.

#### Assessing bias and variance

For a set of parameters, the estimates for MOI parameter and haplotype frequencies were obtained for each of the *S* datasets 𝒳^(1)^, …, 𝒳^(*S*)^, yielding *S* estimates 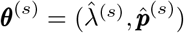, *s* = 1, …, *S*. Furthermore the mean MOI was calculated using

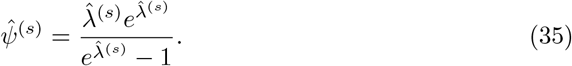

The quality of the estimators was quantified using precision and accuracy. Depending on the parameter scale, these were assessed using either absolute or relative measures. Let *θ* be the parameter of interest, and 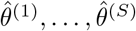 the estimates of the *S* generated datasets. The relative bias of the estimator 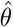 is empirically estimated as

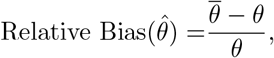

where 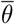 denotes the empirical mean

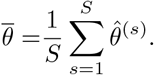

The coefficient of variation (CV) was used to measure variability relative to the true parameter value as

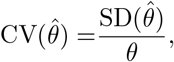

where the empirical standard deviation is given by

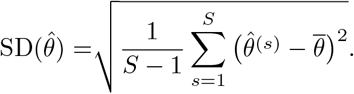

Furthermore, to assess whether the empirical precision of the MLE agrees with theoretical expectations, the empirical standard deviations were compared with the asymptotic standard deviations derived from the Fisher Information Matrix. This comparison tests whether the estimator’s observed variability aligns with the Cramér-Rao lower bound, as derived in section Fisher Information and the Cramér-Rao Lower Bound.

All numerical simulations were implemented in R version 4.5.1. The code is available at GitHub https://github.com/Maths-against-Malaria/Estimating-HRP-deletions.

## Ethics statement

The study was approved by the Institutional Ethics Committees of the ICMR-National Institute of Research in Tribal Health (NIRTH), Jabalpur, Madhya Pradesh, and the ICMR-National Institute of Malaria Research (NIMR), New Delhi, India (IRB number 201901/29). Written informed consent was obtained from all adult participants prior to sample collection. For participants under 18 years of age, written informed consent was obtained from their parents or legal guardians.

## Empirical data

To demonstrate the application of the method, data from two previously published studies, approved as described above, were analyzed. The first study (“hospital study”) was carried out at Late Baliram Kashyap Memorial Medical College (formerly Maharani Medical College and Hospital) from July 2020 to November 2022 and enrolled malaria positive patients. The second (“community study”) was an active surveillance study in a tribal community, conducted at Darbha Community Health Centre, Jagdalpur, Chhattisgarh, India and included data collected from febrile community members between October and November 2021.

The studies and molecular techniques were described in detail in [47]. In both studies, samples were screened for deletions in *Pfhrp*2 and *Pfhrp*3. In brief, Haplotypes were characterized using eight regions across *Pfhrp*2 and *Pfhrp*3. For each gene, four regions were sequenced: the upstream 5’ region, which consists of non-coding DNA immediately preceding the gene; exon 1–2, which encompasses the intron region between exons 1 and 2; exon 2, which contains the major protein coding region responsible for HRP2/HRP3 antigen expression; and the downstream 3’ region flanking exon 2, which consists of non-coding DNA. These markers enable distinction between complete gene deletions, in which all markers indicate the gene is missing, and partial deletions, in which only specific markers show deletions. Additionally, one neutral marker each at *Pfmsp*1 (encoding *P. falciparum* merozoite surface protein 1) and *Pfmsp*2 (encoding *P. falciparum* merozoite surface protein 2) were included. These highly polymorphic surface protein genes serve as neutral markers for estimating multiplicity of infection and distinguishing between genetically distinct parasite strains within mixed infections. The datasets are available as supplementary material in S5 Empirical data.

## Data availability

All empirical datasets used in this study are provided as Supporting Information files (S5 Empirical data). The R code used to analyze these data is also provided as Supporting Information (S2 R Script). In addition, the R code and empirical datasets will be made publicly available on GitHub at https://github.com/Maths-against-Malaria/Estimating-HRP-deletions.

## Supplementary Materials

**S1 Appendix**

**S2 R Script**

**S3 User manual for the R implementation**

**S4 Example data**

**S5 Empirical data**

## Acknowledgments

The first author sincerely thanks Profesor Alois Pichler (TU Chemnitz) for his supervision, valuable guidance, and continuous support throughout this research. The authors gratefully acknowledge the African Institute for Mathematical Sciences (AIMS) Cameroon for fostering the collaboration between the authors and supporting this research.

The authors also appreciate the many fruitful discussions with friends and colleagues that helped to improve the manuscript.

